# Cell-type specific optogenetic fMRI on basal forebrain reveals functional network basis of behavioral preference

**DOI:** 10.1101/2023.07.06.548057

**Authors:** Yijuan Zou, Chuanjun Tong, Wanling Peng, Yue Qiu, Jiangxue Li, Ying Xia, Mengchao Pei, Kaiwei Zhang, Weishuai Li, Min Xu, Zhifeng Liang

**Author notes:** Correspondence should be addressed to Min Xu or Zhifeng Liang. These authors contributed equally (Yijuan Zou, Chuanjun Tong).

## Abstract

The basal forebrain (BF) is a complex structure that plays key roles in regulating various brain functions. However, it remains unclear how cholinergic and non-cholinergic BF neurons modulate large-scale functional networks and their relevance in intrinsic and extrinsic behaviors. With optimized awake mouse optogenetic fMRI approach, we revealed optogenetic stimulations of four BF neuron types evoked distinct cell-type specific whole-brain BOLD activations, which could attribute to BF-originated low dimensional structural networks. Additionally, optogenetic activation of VGLUT2, ChAT and PV neurons in BF modulated the preference of locomotion, exploration and grooming, respectively. Furthermore, we uncovered the functional network basis of the above BF-modulated behavioral preference, through a decoding model linking the BF-modulated BOLD activations, low dimensional structural networks, and behavioral preference. To summarize, we decoded functional network basis of differential behavioral preference with cell-type specific optogenetic fMRI on BF, and provided an avenue for investigating mouse behaviors from a whole-brain view.

## INTRODUCTION

The basal forebrain (BF) is a set of heterogeneous telencephalic structures on the medial and ventral parts of the cerebral hemispheres^1,2^. The basal forebrain is most commonly known as a key cholinergic brain region^1–3^. However, cholinergic neurons only represent 10–20% of all BF neurons, and many more non-cholinergic neurons, including glutamatergic, parvalbumin-positive (PV) and somatostatin-positive (SOM) GABAergic neurons, are present in BF^1–3^. Previous studies reported that the above four types of BF neurons exhibit both complex local connectivity and long-range projections to the cortex^3,4^. Traditionally, studies of BF have focused on its cholinergic neurons and revealed its involvement in a variety of brain functions, including arousal, attention, memory, learning, motivation and reward both in health and disease^5–12^. However, BF non-cholinergic neurons were also found to contribute to many brain functions^3,13–17^.

As the cholinergic and non-cholinergic BF neurons have divergent long-range projections across mouse brain^4^, it is long speculated BF may contribute to the whole brain functional network dynamics. One recent human fMRI study found global BOLD signal peaks coincided with negative activations in BF, which was associated with arousal fluctuations^18^. Another macaque fMRI study showed unilateral inactivation of BF with muscimol resulted in a decrease in the amplitude of BOLD signal in ipsilateral regions, which receive strong projection from BF neurons^19^. Both studies suggested that the extensive variations in large-scale cortical activity observed through fMRI were linked to BF, which was assumed to be contributed by BF cholinergic neurons. However, in those two studies, direct evidence was lacking to attribute such modulation to BF cholinergic system.

Specific modulation of the cholinergic system, combined with rodent fMRI, revealed its impact on large-scale network activity^20–22^. Chemogenetic activation of BF cholinergic neurons^22^ or pharmacological inhibition of the cholinergic system^20,21^ in anesthetized rats, both demonstrated significant decreases in functional connectivity (FC) and BOLD amplitude within the default mode-like (DMN-like) network, which appears contradictory. Neurophysiological studies in rats showed gamma oscillations were synchronized and of high gamma power between the BF and DMN-like nodes (i.e., anterior cingulate area (ACA), retrosplenial cortex (RSP)) during internally oriented behaviors, such as quiet wakefulness and grooming^23,24^. Furthermore, optogenetic activation of BF PV neurons enhanced BF gamma oscillations and promoted grooming behavior^25^, suggesting the regulation of non-cholinergic neurons to DMN-like network and internally oriented behavior. However, It remains unclear the relationship among the cell-type specific BF modulations, large-scale functional networks and their behavioral relevance.

Therefore, in this study, we established a decoding model to link the BF-modulated large-scale BOLD activations and corresponding behavioral preference via combining (1) the cell-type specific opto-fMRI on BF VGLUT2, ChAT, PV and SOM neurons, (2) the BF cell-type specific anatomical projections and (3) optogenetic free-moving behavioral test. Firstly, through our systematically optimized awake mouse opto-fMRI setup, we found distinct global BOLD responses evoked by the cell-type specific activations of BF neurons. Then, combining the BF cell-type specific anatomical projections, we found BF-originated low dimensional structural networks significantly contributed to cell-type specific opto-fMRI activations. Furthermore, we revealed that activation of BF excitatory (VGLUT2 and ChAT) or inhibitory (PV and SOM) neurons induced externally or internally oriented behavioral preference, respectively. Importantly, we developed a model to link the BF-modulated BOLD activations inside the magnet and behavioral preference outside the magnet, and decoded the global patterns of internally and externally oriented behaviors. In conclusion, we revealed both cholinergic and non-cholinergic neurons modulated global functional networks with corresponding internally and externally oriented behavioral preferences, and provided a new perspective to link the BOLD activations to behavioral relevance.

## RESULTS

### Optogenetic stimulations of BF neurons evoke distinct cell-type specific BOLD activations

We combined Cre-dependent virus (AAV2/9-hSyn-DIO-ChrimsomR-mCheery-WPRE-pA) and VGLUT2-, ChAT-, PV- and SOM-Cre mice to express of ChrimsomR in glutamatergic, cholinergic, PV and SOM GABAergic neurons in BF (Figure 1A and S1). To accommodate the limited space in cryogenic coil, flexible PMMA optical fibers were used in combination with our previous awake mouse fMRI^26–28^ to establish the awake mouse opto-fMRI setup (Figure 1A-D). The biband EPI pulse sequence^28^ was employed to achieve faster acquisition (TR = 500 ms) while maintaining whole brain coverage (in-plane resolution = 150 μm^2^ with 38 axial slices).

**Figure 1.**
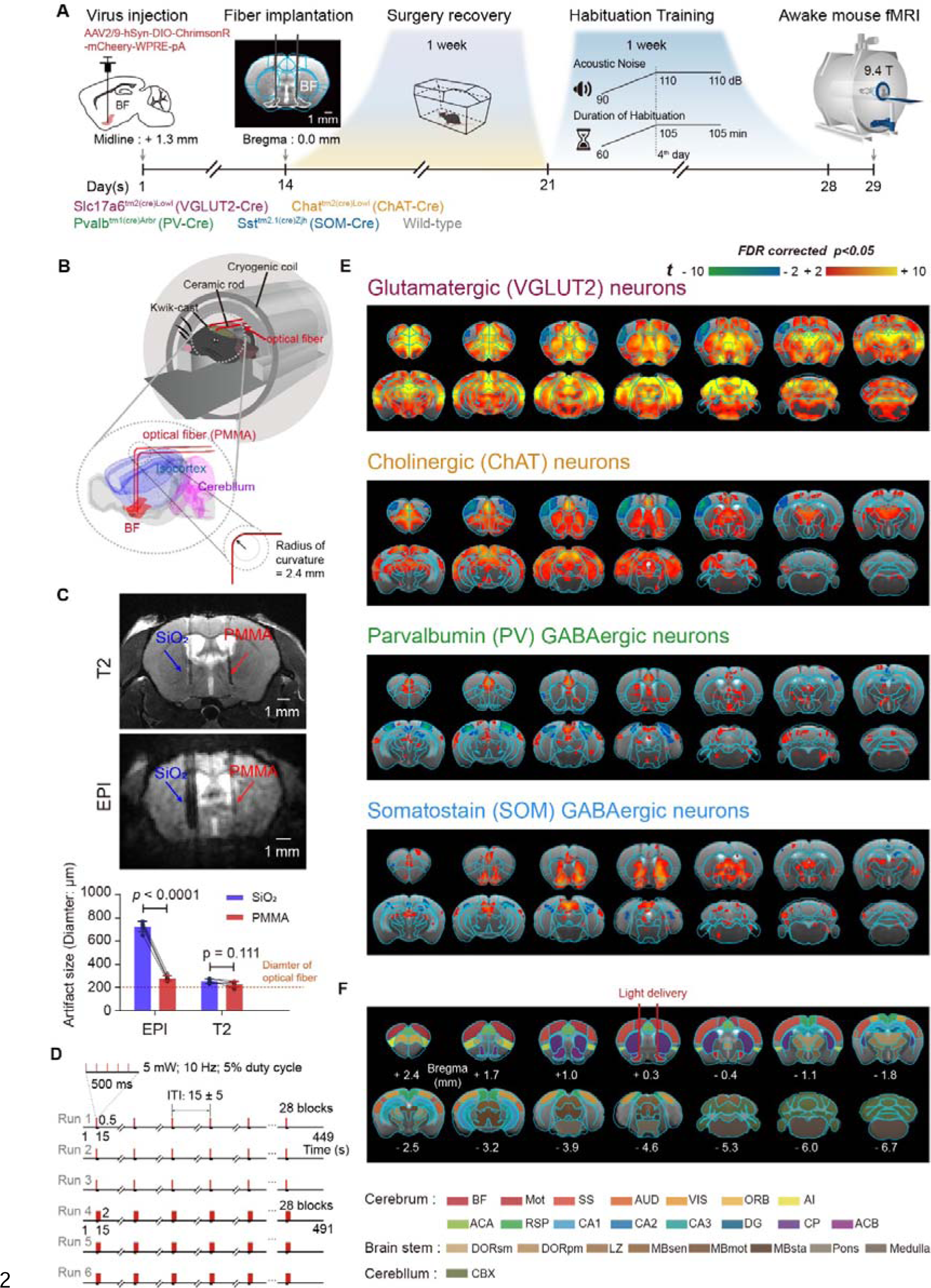
Whole brain opto-fMRI responses driven by cell-type specific neuronal activations of the basal forebrain in awake mice. (A) Schematic setup of opto-fMRI and experimental timeline. (B) Flexible PMMA fiber for optogenetic light delivery within cryogenic coil. Light was delivered to bilateral basal forebrain (BF) via two PMMA optical fibers. PMMA, polymethylmethacrylate. (C) Reduced image artifact by PMMA optical fiber compared to SiO2 fiber. SiO2, silicon dioxide. Statistical significance was determined using paired t-test (two tails, N=5 animals). Error bar, the standard error of the mean. (D) Optogenetic stimulation paradigm. (E) BOLD responses under optogenetic stimulations of cell-type specific BF neurons with FDR corrected p < 0.05. (F) Region-of-interest (ROI) definitions based on CCFv3 Allen mouse brain atlas. Abbreviations were listed in Supplementary Table 1. See also Figure S1-6 and Table S1.

Using our optimized awake mouse opto-fMRI setup and standardized data processing protocol (Figure1A and Figure S2)^26–29^, we observed robust and cell-type specific large-scale BOLD fMRI activations (Figure 1E-F, Figure S3-6). Considering the high spatial similarities between 0.5-s duration and 2-s duration for four BF cell-types (Figure S3), we chose to pool EPI runs with 0.5-s and 2-s durations together, and only focused on the cell-type specific activations (Figure 1E-F). Activation of VGLUT2 neurons in the BF elicited the strongest and most widespread BOLD activations throughout the brain among all four cell types (Figure 1E, upper panel). Activation of ChAT neurons resulted in positive activations across extensive cortical and subcortical regions, including the prefrontal cortex (PFC), ACA, RSP, orbital (ORB), visual (VIS), auditory (AUD), entorhinal (ENT) areas as well as a large proportion of striatum and thalamus (Figure 1E, upper middle panel). In PV-Cre group, ACA showed significant positive activation (Figure1E, lower middle panel). In SOM-Cre group, significant positive activations were observed in ACA, striatum and anterior thalamus (Figure 1E, lower panel). Interestingly, the VGLUT2- and ChAT-Cre groups exhibited significant negative activation constrain within the motor (Mot) and somatosensory (SS) areas.

Notably, we consistently observed positive activations in the DMN-like regions (Figure1E and Figure 2A-C). Importantly, BF PV neurons exhibited the highest specificity for DMN-like activations (Figure 2D and Figure S7) rather than VGLUT2 neurons. Such a phenomenon was consistent with the previous finding in rats that optogenetic stimulation of BF PV neurons activated the DMN-like network and associated behaviors^25^.

**Figure 2.**
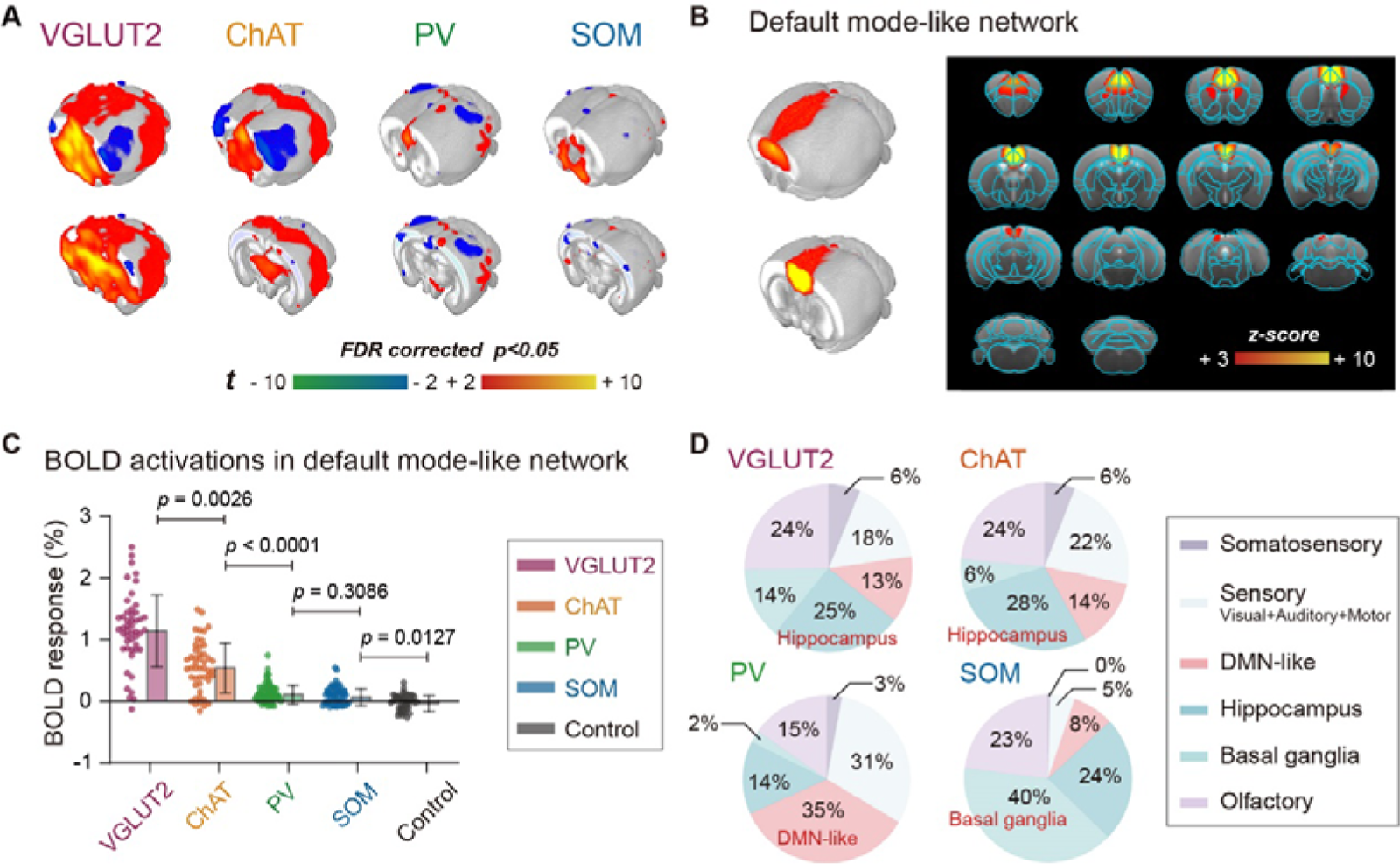
Consistent activations in mouse default mode-like network (DMN-like) under cell-type specific optogenetic stimulations of basal forebrain neurons. (A) BOLD activation maps in 3D view under cell-type specific optogenetic stimulations of basal forebrain (BF) neurons. The activation maps were same as Figure 1E. (B) DMN-like component of mouse in 3D view (left panel) and axial slice view (right panel). (C) Significant BOLD activations in DMN-like regions under cell-type specific manipulation on BF neurons. Statistical significance was determined using a one-way ANOVA followed by Tukey’s post hoc test for group comparison. Each dot represented an individual EPI run. Error bar, the standard deviation (S.D.) of the mean. (D) Highest specificity of DMN-like activations modulated by BF PV neurons. Pie charts showing the composition by mouse functional networks for the cell-type specific opto-fMRI responses. See also Figure S7.

### BF-originated low dimensional structural networks significantly contribute to cell-type specific opto-fMRI activations

To investigate whether the functional activation was directly governed by corresponding anatomical projections from BF neurons, we adopted the previously published anterograde axon tracing data^4^ and registered to our standard MRI space (Figure 3A and 3B). Cholinergic BF neurons displayed widely spread projections to the cortex, while the non-cholinergic neurons mainly innervated subcortical brain regions and had limited projections to most parts of the cortex (Figure 3B). To evaluate the structural-functional correspondence across BF cell types, we calculated the spatial correlation between functional activations (Figure 1E) and anatomical outputs (Figure 3B). Weak structural-functional correspondence was found for all BF cell types (Figure 3C, brown arrows and Figure S8), suggesting direct anatomical projections weakly contributed to opto-fMRI activation patterns. Additionally, we verified that anatomical tracing using the optogenetic protein ChrimsonR-mCherry was consistent with our previously published data^4^ (Figure S9), thus did not impact our results.

**Figure 3.**
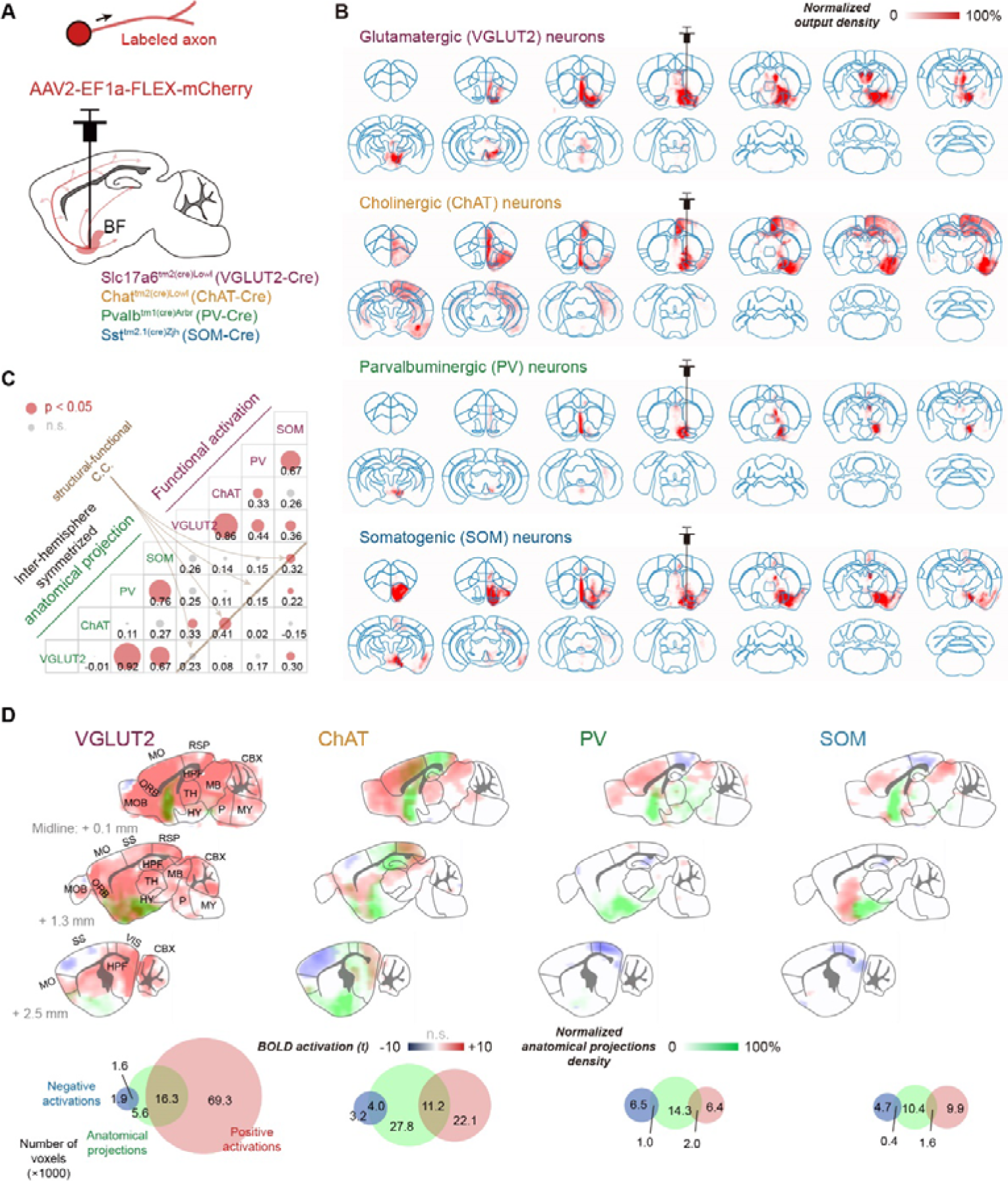
Direct anatomical projections weakly contribute to opto-fMRI activation patterns of BF neurons. (A) Viral injection procedure for tracing BF projections. (B) Whole-brain anatomical projections from 4 BF neuron types. (C) Weak spatial similarities (brown line) between anatomical projection and functional activation maps for four BF neuron types. The size of each circle indicates the spatial similarity (Pearson’s correlation coefficients, C.C.) between the opto-fMRI activations and anatomical projection density across 213 brain regions. The parcels of these 213 brain regions were chosen based on the Allen Mouse Brain Connectivity Atlas, covering all voxels across mouse brain. No thresholds were used to select pixels. (D) Spatial maps (upper panel) and quantitative evaluations (lower panel) of the overlay between anatomical projections and functional activations. See also Figure S8-9.

One potential contributor to the functional responses was secondary projections of the regions that receive direct projections from BF neurons (Figure 4A). To examine such a hypothesis, we generated the cell-type specific BF secondary projection matrix by spatial weighting the voxel-wise whole-brain mesoscopic connectivity matrix^30,31^ based on the BF direct projections (Figure 4B & Figure S10A-C). We found two-synaptic connections, i.e., 1^st^ × 2^nd^ anatomical connections, showed higher spatial similarity of cell-type specific opto-fMRI activations, compared to direct projections (Figure S11), suggesting that the secondary projections were important contributors for BF four neuron types. Moreover, we concatenated secondary projections from four types of BF neurons to cover all projections originating from BF, including those from local interaction of BF cell types. Then, the concatenated secondary projections were further decomposed using the non-negative matrix factorization (NMF) analysis (Figure 4B & Figure S10A-C). One significant advantage of NMF over other dimensionality reduction techniques is its ability to ensure that all elements in the decomposed low-rank matrix are non-negative, which is crucial for interpreting both NMF components and associated weights as actual network activity.

**Figure 4.**
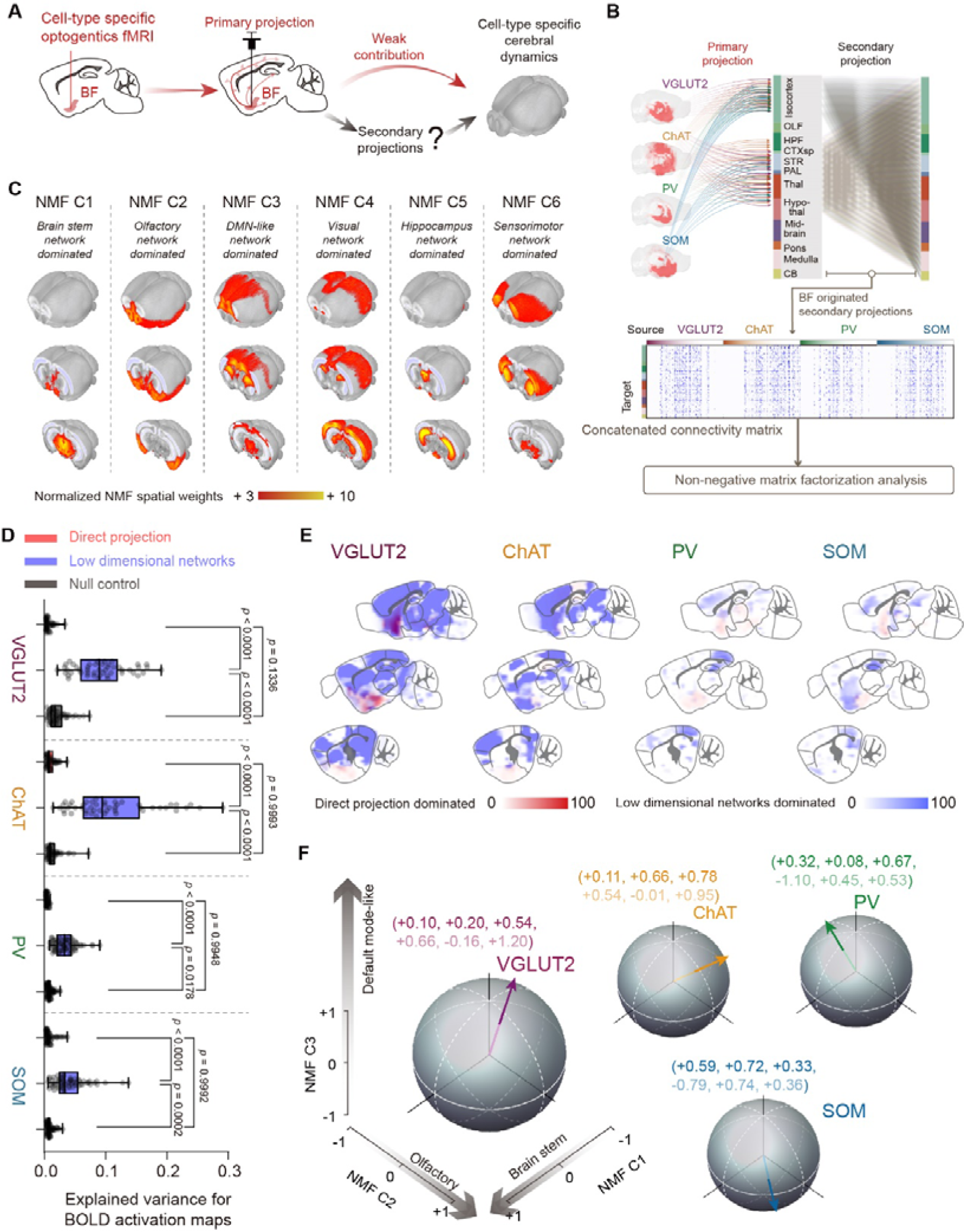
BF originated low dimensional structural networks significantly contributed to cell-type specific opto-fMRI activations. (A) Hypothesis of the contribution of secondary projections on cell-type specific opto-fMRI activations. (B) Computational pipeline of non-negative matrix factorization (NMF) on BF originated whole-brain mesoscopic secondary projections. (C) Spatial maps of three low dimensional structural networks (i.e., the first six NMF components) in 3D view. (D-E) Quantification (D) and spatial distributions (E) of explained variance for opto-fMRI activations from the direct projections and low dimensional structural networks (as in Figure 4C) respectively. Statistical significance was calculated by two-tailed paired t-test. Notably, the null control was constructed by spatially shuffling the BOLD activation maps. Each dot represented an individual EPI run. Box, 25-75% range and median line; whisker, minimum to maximum range; plus mark, the mean value. (F) Cell-type specific opto-fMRI activations in six dimensional NMF space. Feature vector for each BF cell type was normalized by its magnitude. See also Figure S10-12.

Subsequently, we identified six low-dimensional NMF components that were predominantly composed of brain stem, olfactory, DMN-like, visual, hippocampus and sensorimotor network (Figure 4C & Figure S10D-E), respectively. To explore whether the low-dimensional structural networks dominated the cerebral dynamics, we calculated the explained variance for the BOLD activation maps based on the multivariate regression. Such regression approach was applied (Figure 4D) between empirical BOLD activations and 1) cell-type specific direct projection or 2) best combinations of BF originated low dimensional networks or 3) the null control of BOLD activation maps, respectively. The null control maps were constructed by spatially shuffling the BOLD activation maps with spatial autocorrelation preserved. We observed that low-dimensional structural networks exhibited significantly higher explained variance in cell-type specific opto-fMRI responses compared to the direct projections and the null control (Figure 4D-E and Figure S12), indicating their dominant effect of 2^nd^-anatomical connections on opto-evoked cerebral dynamics. The scaling coefficients of NMF components were then defined as optimal feature vectors, representing the engagement levels of NMF components under optogenetic activation of BF neurons (Figure 4F). Therefore, these findings provide evidence that BF originated low dimensional structural networks significantly contributed to cell-type specific opto-fMRI activations.

### Optogenetic activation of VGLUT2, ChAT and PV neurons in BF modulated the preference of locomotion, exploration and grooming, respectively

We further investigate whether cell-type specific modulation of BF exerted an impact on mouse internally and externally oriented behaviors. The behavioral test was conducted in the arena under the same optogenetic stimulation paradigms as in the fMRI experiments to examine the locomotion speed, quiet wakefulness, grooming and object explorations (Figure 5A-C). The VGLUT2-Cre group exhibited a significantly higher locomotion speed (Figure 5D), which was consistent with the wakefulness and avoidance behavior-promoting effect of BF VGLUT2 neurons^3,15,16^. The ChAT-Cre group demonstrated significantly higher novel object explorations (Figure 5E), which is consistent with the involvement of cholinergic systems in attention, learning, and memory^11,12^. There were no discernible differences in familiar object exploration among the groups (Figure S13A). Moreover, optogenetic stimulation of BF PV neurons resulted in the highest DMN-associated behavior, specifically grooming (Figure 5G), which is consistent with the previous study in rats^25^. SOM-Cre group exhibited a greater duration of quiet wakefulness in comparison to other subpopulations (Figure 5F).

**Figure 5.**
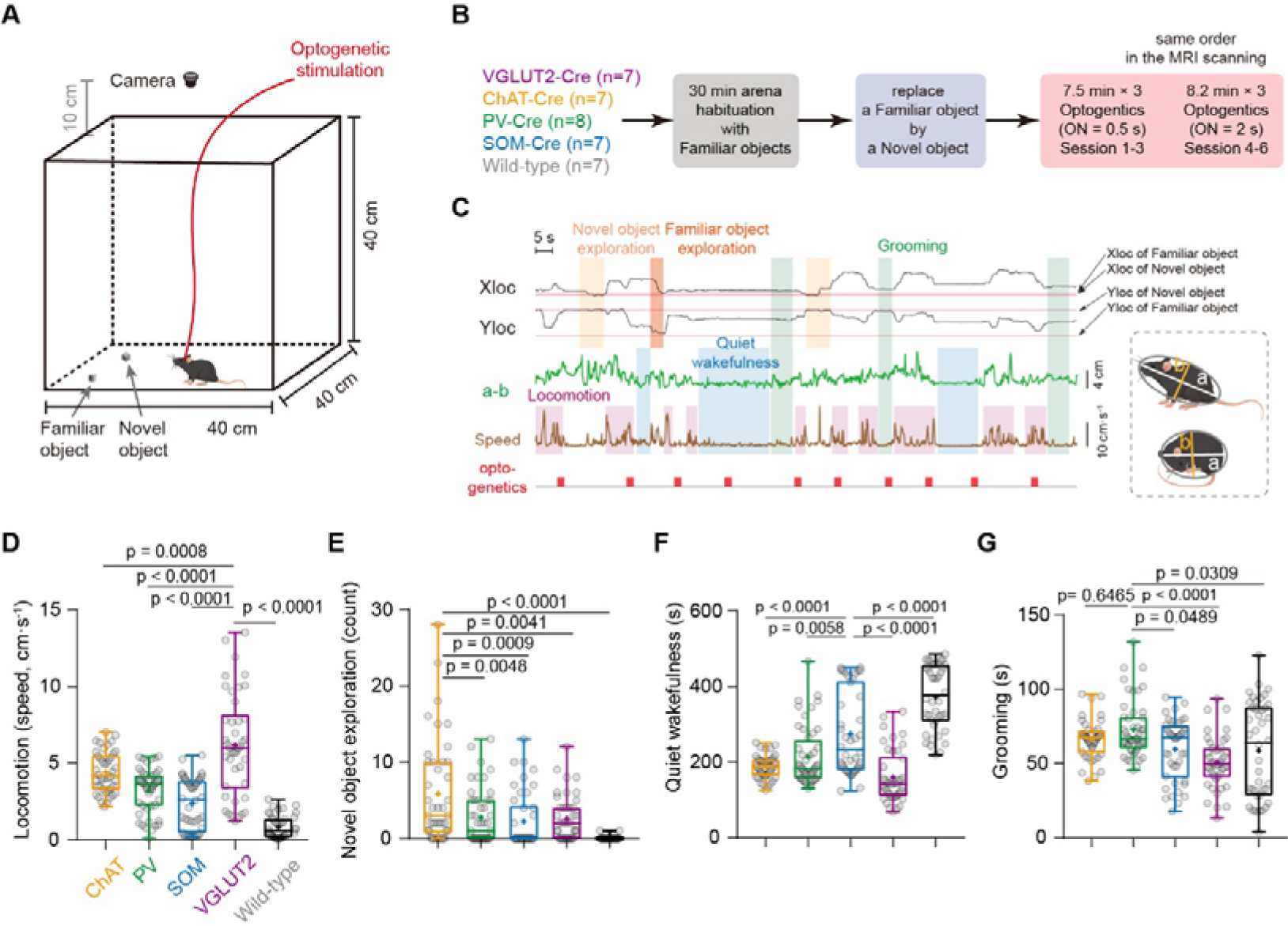
Optogenetic activation of VGLUT2, ChAT and PV neurons in BF modulated the preference of locomotion, exploration and grooming, respectively. (A-B) Schematic of the free-moving behavioral experiments under optogenetic stimulations. (C) Representative behavioral traces. (D-G) Behavior performances under optogenetic activations of four BF neuron types, including locomotion (D), novel object exploration (E), quiet wakefulness (F) and grooming (G). Details of multiple comparison across cell-types were list in supplementary Table 2. Each dot represented the averaged behavioral performance in each experimental session. Box, 25-75% range and median line; whisker, minimum to maximum range; plus mark, the mean value. See also Figure S13-15 and Table S2.

Additionally, behavioral preference modulated by BF neuron types remained consistent throughout all six runs and was primarily driven by distinct cell types rather than other factors such as duration and experimental runs (Figure S13B-C). Furthermore, we examined whether the difference between genetically modified Cre-mice and wild-type mice potentially influenced the behavioral performance (Figure S14). The highly similar results of BOLD activations (Figure S14A-B) and behaviors (Figure S14C) between the PV-cre control group and wild-type control group indicated that using wild-type mice with the same optogenetic stimulation paradigm as the control group was less likely to influence our conclusions.

Next, we sought to explore why the locomotion and BOLD activations appeared incoherent in sensorimotor regions. We used the calcium fiber photometry to record opto-evoked neuronal calcium activities in SSp-m (the negatively activated region) and RSP (the positively activated region) for VGLUT2 and Control group (Figure S15A). We found optogenetic activations of BF VGLUT2 neurons evoked significant elevations of calcium activities in RSP and SSp-m (Figure S15B-C) compared to those in the control group. We inferred that the initial rise of BOLD signals in SSp-m (Figure S4-5) and higher locomotion speed (Figure 5D) in the VGLUT2 group were associated with enhanced neural calcium activity. Therefore, the initial positive BOLD responses, locomotion increase and calcium responses were aligned with each other in sensorimotor regions. Furthermore, we speculated that the subsequent decrease of BOLD responses (Figure S4-5) might relate to the arousal elevation evoked by the activation of BF VGLUT2 and ChAT neurons^18,19,32,33^.

### Decoding model links BF-modulated BOLD activations and internally and externally oriented behavioral preference

As the same optogenetic stimulation paradigm was applied in opto-fMRI and behavioral test, we hypothesized that cell-type-specific activation maps and behavior-related cerebral patterns share common low-dimensional (NMF) space (Figure 6A). Therefore, we constructed a decoding model (Figure 6B & S16) to predict the behavior related global patterns, based on the feature vectors of BOLD activations (Figure 6C) and behavioral performance (Figure 5D-G). The model’s regression coefficients were identified as behavior-related feature vectors, representing predicted cerebral patterns of each behavior in the NMF space (Figure 6D). The minimal deviation angle (Figure 6E, red shadow) between feature vectors of BOLD activations (Figure 6C) and behaviors (Figure 6D) indicated the behavioral preference, e.g., the feature vector of VGLUT2 group had minimal deviation angle with that of locomotion (Figure 6E), corresponding to its preference on locomotion (Figure 5D). The significantly negative correlations between the deviation angles and empirical behavioral results (Figure 6F) further suggested that low dimensional network organization supported the behavioral preference of BF cell types.

**Figure 6.**
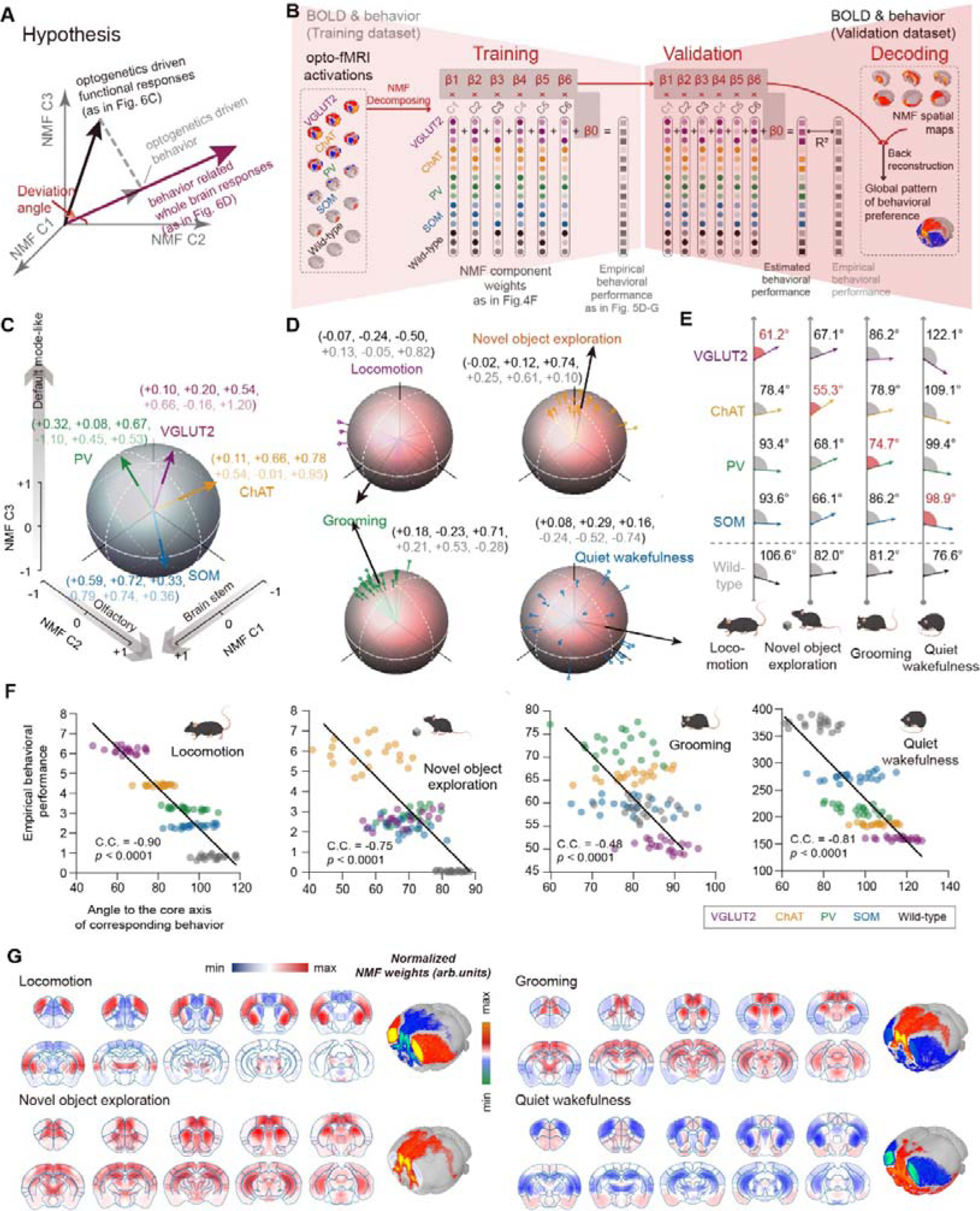
A decoding model predicted the global patterns of opto-induced behaviors outside the magnet based on the low dimensional networks. (A) Hypothesis: cell-type specific activation maps and behavior-related cerebral patterns shared common low-dimensional (NMF) space as the same optogenetic stimulation paradigm was applied in opto-fMRI and behavioral test. (B) Schematics of decoding model to link BF modulated BOLD activations and behavioral preference. Details were shown in Figure S20. (C-D) Feature vectors of cell-type specific BF modulations (C, as in Figure 3G) and predicted behavior-related cerebral patterns (D). Colored lines and black arrows, predicted and corresponding averaged feature vectors for each behavior. Each colored line represented an individual decoding process (N = 20). We randomly chosen 3 sessions as training dataset out of 6 sessions for each process and repeated this process to cover all combinations. (E) Deviation angles between feature vectors in Figure 6C and Figure 6D. Red shadow represented the minimal angle across cell-type specific BF modulations in each behavior (wild-type excluded), indicated the most distinct behavioral preference. (F) Significant correlations between deviation angles and empirical behavioral performance. Smaller deviation angles indicated shorter distances to the behavioral feature vectors in NMF space, i.e., higher behavioral preference of BF cell types. Each dot represented an individual decoding process (N = 20, as in Figure 5D-G). Black line represented the best linear fit. C.C., Pearson’s correlation coefficients. (G) Predicted cerebral responses of mouse behaviors based on the low dimensional NMF space. See also Figure S16.

Based on the behavior-related feature vectors (Figure 6D) and the spatial patterns of NMF components (Figure 3C), we further predicted the cerebral responses of behaviors (Figure 6B, right panel and Figure 6G). Interestingly, the predicted global pattern of grooming showed opposing patterns between intrinsic networks (i.e., DMN-like network) and extrinsic networks (e.g., salience network) (Figure 6G, right panel), which was consistent with previous findings in rats^34^ and human^35–38^. The predicted global pattern of quiet wakefulness (Figure 6G, right panel) exhibited high spatial similarity to that of grooming, and both of these were opposing to that of locomotion (Figure 6G, left panel). Moreover, the predicted global pattern of novel exploration (Figure 6G, left panel) exhibited robust positive responses in the prefrontal cortex and hippocampus, consistent with previous evidence of cholinergic innervation to these regions involved in novelty preference^39–41^.

To verify the predicted spatial patterns of mouse behaviors, we first conducted the awake mouse fMRI scanning with simultaneous behavioral monitoring using two customized MR-compatible cameras (Figure 7A). We observed apparent grooming behavior during the scanning (Figure 7B) and corresponding functional activations (Figure 7C). Significant positive correlation between the empirical (Figure 7C) and predicted (Figure 6G) spatial maps indicated the validity of predicted global patterns of mouse behaviors (Figure 7D and Figure S17).

**Figure 7.**
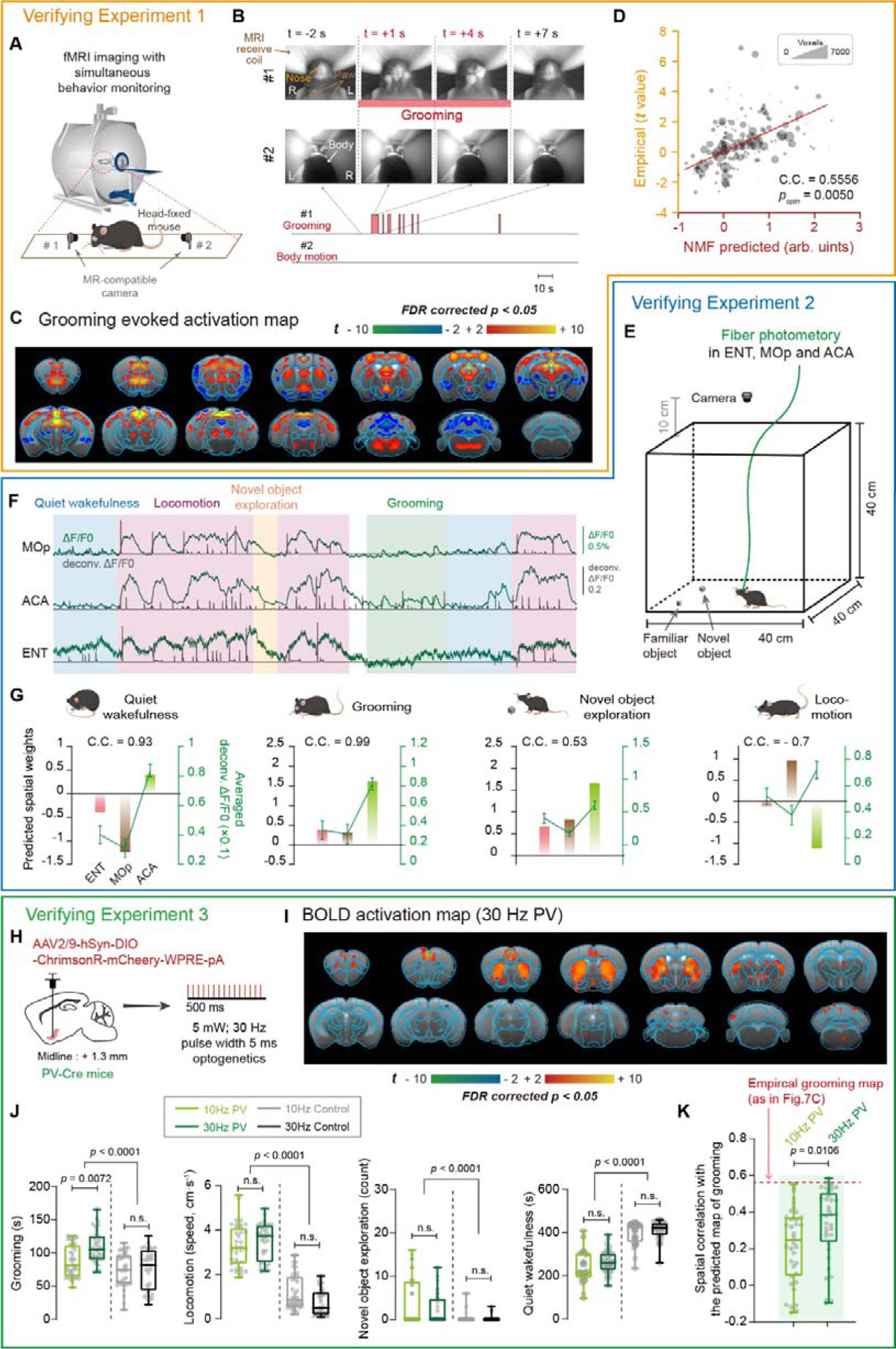
Experimental validations of predicted behavior-related cerebral responses. (A) Schematic of the fMRI experimental setup with simultaneous behavior monitoring. (B) Example of spontaneous grooming during awake mouse fMRI scanning. (C) Empirical activation maps of mouse spontaneous grooming. (D) Significant correlation between empirical and predicted activation maps of mouse grooming. Each dot represented an individual brain region, scaled according to the voxel number of each region. C.C., Pearson’s correlation coefficients. Significance test was determined using the spatial autocorrelation preserving shuffling (E) Schematic of the free-moving behavioral experiments with simultaneous calcium fiber photometry in primary motor area (MOp), anterior cingulate area (ACA) and entorhinal cortex (ENT) respectively. (F) Representative calcium signals in MOp, ACA and ENT regions during spontaneous behaviors. (G) High correlation between the predicted responses and empirical neuronal activity in MOp, ACA and ENT during quiet wakefulness, grooming and novel object exploration, except for locomotion. C.C., Pearson’s correlation coefficients. Error bar, the standard error of the mean. (H) Schematic diagram depicting fMRI scanning and behavioral test under the 10-Hz and 30-Hz optogenetics on BF PV neurons. All experimental conditions were same as Main Figure 1 and 5. (I) BOLD activations map of 30-Hz optogenetics on BF PV neurons. (J) 30-Hz optogenetics on BF PV neurons evoked more grooming behavior for mice than that of 10-Hz optogenetics. Statistical significance was determined using the one-way ANOVA with Tukey’s multiple comparison test. Each dot represented an individual experimental session. Box, 25-75% range and median line; whisker, minimum to maximum range. (K) Activation map of 30-Hz optogenetics on BF PV neurons showed higher spatial similarity to the predicted grooming pattern, compared to those of 10-Hz. Statistical significance was determined using the two-tailed two sample t-test. Each dot represented an individual EPI run. See also Figure S17.

We further conducted another verifying experiment with simultaneous calcium fiber photometry in ACA, primary motor area (MOp), and entorhinal cortex (ENT) during free-moving behaviors (Figure 7E-F). We found a notable positive correlation between the predicted spatial weights and neural activities (Figure 7G) during quiet wakefulness, grooming, and novel object exploration. The negative correlation during locomotion was unexpected but somewhat reasonable, as the locomotion consistently elicits widespread activation and can potentially confound the analysis of other behaviors^42–45^. Thus, the above two verifying experiments highlighted the reliability of our decoding model and the resulting predicted global patterns of mouse behaviors. Therefore, our results provided a framework for behavioral research through the lens of cerebral network basis, linking the BF-modulated BOLD activations to the internally and externally oriented behavioral preference.

Moreover, we conducted additional experiments with 30-Hz optogenetic stimulation on BF PV neurons as PV neuron is characterized by its fast spiking activity^46^. All other conditions were kept the same but with a higher frequency 30 Hz (Figure 7H). The results revealed 30 Hz optogenetic stimulation elicited significant increased BOLD activations in DMN-like regions (Figure 7I) as well as increased DMN-associated behaviors such as grooming (Figure 7J), compared to those from 10 Hz. Importantly, activation map of 30-Hz optogenetics on BF PV neurons showed higher spatial similarity to the predicted grooming pattern compared to those of 10 Hz (Figure 7K), providing further support for the reliability of our decoding model for linking the BOLD activation maps and mouse behaviors.

## DISCUSSION

The functional roles of BF cholinergic and non-cholinergic neurons, from large-scale functional networks to behaviors, remain elusive. We first demonstrated cell-type specific whole-brain responses and corresponding internally and externally oriented behavioral preference induced by optogenetic activation of four BF neuron types. Subsequently, we developed a decoding model to link the BF-modulated BOLD activations and behavioral preference based on the organization of a low-dimensional cerebral network. Ultimately, we predicted the global patterns of mouse behavioral preference via our model, which were verified by two different experiments, i.e., (1) awake mouse fMRI with simultaneous grooming monitoring and (2) calcium fiber photometry during free-moving behavior test. In summary, our results demonstrated that both cholinergic and non-cholinergic BF neurons modulate cerebral functional networks and corresponding internally and externally oriented behavioral preference. Our results also provided a framework for investigating animal’s behaviors in the whole brain view.

It is not trivial to acquire high-quality opto-fMRI data in mice, with many limiting factors such as anesthesia, low SNR, and image artifacts caused by fiber implants. So far, most previous opto-fMRI studies in rodents have been conducted under anesthesia^47–54^, with only a few exceptions^55–57^. To avoid the confounding factors of various anesthesia methods^58,59^, we established the awake opto-fMRI setup based on our extensive experience in awake mouse fMRI^26–29^. Furthermore, a multiband EPI pulse sequence was employed to acquire functional images to achieve faster acquisition while maintaining whole brain coverage^28^. To take advantage of the high SNR provided by the cryogenic RF (radio frequency) coil, a flexible PMMA optic fiber implant was used to accommodate the limited space of the cryogenic coil (Figure 1B). Importantly, PMMA has similar magnetic susceptibility to that of the brain tissue^60^ thus reduced imaging artifacts caused by fiber implantation (Figure 1C). In conclusion, we have optimized the setup for awake mice opto-fMRI systematically to acquire high-quality opto-fMRI data.

With the optimized approach described above, we revealed cell-type specific whole-brain responses under optogenetic activation of BF neurons. The most strong and widespread BOLD responses (Figure 1E, upper panel) and corresponding highest locomotion speed (Figure 5D) of BF VGLUT2 group possibly attribute to the promoting wakefulness and avoidance behavior effects of BF VGLUT2 neurons^3,14,15^. The activation of the PFC, Mot, SS, AUD, and striatum (Figure 1E, upper middle panel) and corresponding highest novel object exploration (Figure 5E) of the ChAT group were consistent with the BF cholinergic neuron involved circuits that modulate recognition memory, sensory perception and attention regulation^39,41,61–64^. The higher quiet wakefulness (Figure 5F) of SOM group may attribute to the sleep-promoting^3^ and locomotion-inhibiting effects^65^ of BF SOM neurons. Importantly, PV group showed positive activation mainly within DMN-like network, resulting the highest specificity for DMN-like activations of PV group than other groups (Figure1E, lower middle panel and Figure 2D), along with the corresponding highest grooming (Figure 5G). The above results were consistent with previous work on BF PV neurons’ modulation on DMN-like networks and internally oriented behaviors^25^. Interestingly, we consistently observed positive activations in the DMN-like regions (Figure 1E and Figure 2) under the activation of all four BF neuron types, indicating the complex yet robust interaction between BF and cortical DMN-like network. It also supports the notion that BF may be part of a subcortical DMN network^24,25,36,66–68^. Therefore, the current study provided a more complete perspective for whole-brain functional responses and corresponding behavioral preference from activations of four BF neuron types.

We noticed that the behavioral differences among BF cell types were not drastic (Figure 5). It can be potentially attribute to 4 reasons: (1) The primary function of BF is to modulate brain states^3,14,69,70^. It is not surprising that BF plays a modulatory role in biasing the probability of these complex behaviors^15,25,39,41,61–64^. Therefore, it is unlikely that the activation of one BF neuron type would elicit a strong one-to-one corresponding behavior; (2) The complex local connections among four BF neuron types^3^. Consequently, optogenetic stimulation of each BF neuron type could elicit responses from other BF cell types, collectively contributing to mouse behaviors; (3) The long familiarization and behavioral test time. Our behavioral test paradigm was not conventional in order to keep consistent with fMRI scanning paradigm and further enable the subsequent linking between BOLD activations and free-moving behaviors. Therefore, we had to use the long familiarization and behavioral test time, during which, mice tended to become bored and entered immobile state. Such long familiarization and behavioral test time could potentially reduce the behavioral impact of optogenetic modulation; (4) Enhanced modulation effects on mouse behaviors may be achieved by using different optogenetic frequency, particularly for PV neurons (Figure 7H-K). Indeed, 30 Hz optogenetic stimulation elicited significant increased BOLD activations in DMN-like regions (Figure 7I) as well as increased DMN-associated behaviors such as grooming (Figure 7J), compared to those from 10 Hz. However, considering the comparing BF four neuron types and controlling for potential artifacts arising from different optogenetic frequencies, utilizing the same frequency of 10 Hz may be a reasonable compromise.

It is well known that the primary anatomical projections are key regulators of functional dynamics. Interestingly, we found weak structural-functional correspondence between BF cell-type specific anatomical projections and functional activations. Such phenomenon was consistent with the finding that functional connectivity originated from dopamine neurons in the ventral tegmental area^51^ and serotonin neurons in the dorsal raphe^47^ were uncoupled with corresponding dopaminergic and serotoninergic neuron projection density, respectively. Our results demonstrated the low dimensional structural networks decomposed from BF-originated secondary projections significantly contributed to the cell-type specific BOLD activations. As previous studies demonstrated that the low dimensional structural networks served as the backbone of functional dynamics and cognitions in human or marmoset brain^71,72^, our finding suggested the dominating effects of low dimensional structural networks on functional activations are markedly consistent across species.

However, a framework to systematically establish the link among the cell-type specific BF modulations, global structural networks, and behavioral preference is lacking. Thus, our study built a decoding model to link the BF-modulated global activation patterns and the behavioral preference in a low dimensional network space (Figure 6). After projecting high dimensional BOLD activations into low dimensional space spanned by structural networks, i.e., NMF components (Figure 4C), the dimension of resulting features, i.e., weights of NMF components (Figure 4F or 6C), were much closer to the dimension of mouse behaviors. Using the low dimensional features as model inputs highly improved the computational efficiency. More importantly, such a strategy enhanced the feasibility of capturing discriminative features from BF-modulated BOLD activation patterns, which were subsequently used to model the corresponding behavioral preference. Also, our BF cell-type specific opto-fMRI, including manipulation on four neuron types instead of one^22^, provided more complete and informative BOLD features as the model inputs, improving the robustness of our decoding model.

Interestingly, our modeling approach provided a general methodology to explore the global patterns of behaviors. Based on the decoding model, we back-reconstructed the global patterns of mouse behaviors, such as quiet wakefulness and grooming (Figure6B, right panel and 6G). Thus, the predicted global patterns offered potential “reference templates”, similar to the “arousal template”^33^, and could be used to infer the behavioral occurrence through the spatial correlation between the template and the empirical imaging data^18,33^. Nevertheless, it is difficult to experimentally verify the predicted global patterns (Figure 6G) in the MRI scanner. Thus, we further modified our awake mouse fMRI setup to allow the animal’s forelimbs to groom during the fMRI scanning (Figure 7B), and obtained the empirical grooming evoked BOLD activations (Figure 7C). The significant correlation between empirical and predicted grooming patterns (Figure 7D) provided the experimental validation for our decoding model. Interestingly, the grooming pattern with activations in DMN regions and de-activations in extrinsic networks (e.g., salience network) was in line with BOLD responses during internally focused cognitive state in the previous human studies^35–38^. Together, these results suggested the reliability of our modeling approach, which could be further used in inferring the global patterns of behaviors in future imaging studies.

Previous human imaging studies^35–38^ revealed significant DMN activation and dorsal attention network suppression under internally focused conditions. Such a phenomenon was in accordance with our empirical groom activations (Figure 7C), suggesting the cross-species consistency of the anti-correlated relationship between intrinsic and extrinsic networks^73^. During the external task condition, the BF and DMN were deactivated together in human fMRI studies^35,36,74^. The BF’s prominent structural connectivity with DMN-like cortical regions has been suggested to parallel its known modulatory projections involving cholinergic, GABAergic, and glutamatergic systems^4^. In our study, we revealed the highest specificity for the DMN activations with corresponding grooming preference under the optogenetic activations of BF PV neurons (Figure 2 and 4). Such results agreed with previous studies that projections of BF PV neurons have been specifically linked to the control of DMN state transitions in rats^25,34^. Therefore, we confirmed the BF PV neurons on the DMN activity during the internally oriented behaviors.

An important limitation in our model was the assumption that the same optogenetic stimulation paradigm induced a similar global response inside and outside the magnet. Naturally, the brain state of the mouse was undoubtedly different in those two environments. However, it is not unreasonable to assume that the strong cerebral responses were more dominated by the optogenetic stimulations than the environment. Moreover, our empirical evidence provided experimental support for the validity of our modeling approach (Figure 7). Additionally, it was somewhat arbitrary or insufficient to select these three regions for calcium fiber photometry to validate the reliability of decoding patterns of quiet wakefulness, grooming, novel object exploration and locomotion. Future investigations should include more regions to provide better validation of our decoding patterns of free-moving behaviors. Another limitation was the lack of recording the natural activity of cell-type specific BF neurons during various behaviors, such as grooming, for better confirmation of different BF neuron types associated with behavioral preferences. Furthermore, the current study only examined the cell-type specific effects of BF neuron activation. Future investigations should incorporate cell-type specific inhibition of BF neurons for a more complete view.

Last but not least, the complex interactions among BF neuron types^3^ were not specifically considered in the current model. As demonstrated in our previous study^3^, there are complex local connections among different cell types within BF. Given that VGLUT2 neurons exhibit excitatory connections with ChAT, PV, and SOM neurons, the activation map of BF VGLUT2 neurons may partly result from these local connections as it resembles a combination of activation maps from the other three neural types. Nevertheless, our concatenated secondary projections encompass all possible outputs originating from BF four neuron types. Thus, our modelling results (Figure 4 & 6) were unlikely to be affected by local interactions among BF neuron types. Further biophysical modeling incorporating these local connections may be conducted in the future.

## STAR METHODS

Detailed methods are provided in the online version of this paper and include the following:

- KEY RESOURCES TABLE
- RESOURCE AVAILABILITY

- Lead Contact
- Materials Availability
- Data and Code Availability
- EXPERIMENTAL MODEL AND STUDY PARTICIPANT DETAILS

- Animals
- METHOD DETAILS

- Stereotaxic surgeries
- Habituation for awake mouse optogenetic fMRI (opto-fMRI)
- Optogenetic fMRI acquisition
- Simultaneous behavioral monitoring and fMRI acquisition
- fMRI data processing
- General linear model of cell-type specific optogenetic stimulations and spontaneous behaviors
- Specificity of DMN-like activations driven by BF neurons
- Anterograde axon tracing from BF neurons
- Voxel-wise whole-brain mesoscopic structural connectivity
- Non-negative matrix factorization
- Optogenetic free-moving behavioral test
- Decoding the global patterns of mouse behaviors
- Histology and microscopy
- Free-moving behavioral tests with simultaneous fiber photometry
- Optogenetics with simultaneous fiber photometry
- Significance test using the spatial autocorrelation preserving shuffling
- Comparing imaging artifacts of PMMA and SiO2 fiber optic implants
- QUANTIFICATION AND STATISTICAL ANALYSIS

## ACKNOWLEDGMENTS

The authors thank Jingjing Ye (iHuman Institute, ShanghaiTech Univ.) and Garth J. Thompson (iHuman Institute, ShanghaiTech Univ.) for the help in fMRI data acquisition, Ziyue Wang (CEBSIT, CAS) for the help in mouse surgery, Mouse Animal Facility of CEBSIT for animal care, and the Brain Science Data Center, Chinese Academy of Sciences for dataset release. This work was supported by the National Science and Technology Innovation 2030 Major Program (2021ZD0202200, 2021ZD0200100 to ZL), National Natural Science Foundation of China (32221003 to MX), CAS Project for Young Scientists in Basic Research (YSBR-071 to MX), Lingang Laboratory & National Key Laboratory of Human Factors Engineering Joint Grant (LG-TKN-202203-01 to MX), China National Postdoctoral Program for Innovative Talents (BX20230383 to CT), China Postdoctoral Science Foundation (2023M743616 to CT).

## AUTHOR CONTRIBUTIONS

Y.Z., C.T., M.X. and Z.L. designed the study; Y.Z., C.T., Y.Q., J.L., Y.X., M.P., K.Z. and W.L. collected the MRI and behavioral data; Y.Z., C.T. and Z.L. preprocessed and organized the MRI and behavioral data; W.P. and M.X. collected and preprocessed the anterograde tracing data; C.T., Y.Z. and Z.L. conducted the decoding modeling. Y.Z., C.T., M.X. and Z.L. wrote the original draft and revised the draft. All authors discussed and approved the manuscript.

## STAR METHODS

### RESOURCE AVAILABILITY

#### Lead Contact

Correspondence and requests for materials should be addressed to the Lead Contact, Zhifeng Liang.

#### Materials Availability

This study did not generate new unique reagents.

#### Data and Code Availability

All data reported in this paper has been deposited at Zenodo DOI: https://doi.org/10.5281/zenodo.10499624.

All original code has been deposited at GitHub and is publicly available. DOI is listed in the key resources table.

Any additional information required to reanalyze the data reported in this paper is available from the lead contact upon request.

### EXPERIMENTAL MODEL AND STUDY PARTICIPANT DETAILS

#### Animals

All animal experiments were approved by the Animal Care and Use Committee of the Institute of Neuroscience, Chinese Academy of Sciences, Shanghai, China. Experiments were conducted with male mice at 8-10 weeks of ages, weighted 18-30 g. Number of mice in each experiment was listed in below table. No statistical methods were used to predetermine sample sizes but our sample size is similar to previous publications^56,76^. Wild-type mice were purchased from institute-approved vendors (Shanghai Laboratory Animal Center, or LingChang Experiment Animal Co., China), and ChAT-IRES-Cre, VGLUT2-IRES-Cre, PV-IRES-Cre and SOM-IRES-Cre mice (Jackson stock #: 006410, 016963, 008069 and 013044 respectively) were obtained from Jackson Laboratory. Mice were group-housed (5–6/cage) under a 12-h light/dark cycle (light on from 7 a.m. to 7 p.m.) with food and water *ad libitum*.

**Table.** Summary of Number of mice used in each experiment.

|  | ChAT-Cre | PV-Cre | SOM-Cre | VGLUT2-Cre | Wild-type |
| --- | --- | --- | --- | --- | --- |
| 10-Hz optogenetics with fMRI scanning | 8 | 11 | 10 | 8 | 7 |
| 10-Hz optogenetics with free-moving behavioral test | 7 | 8 | 7 | 7 | 7 |
| 10-Hz optogenetics with fiber photometry in SSp-m & RSP | - | - | - | 8 | - |
| Verifying experiment 1: fMRI scanning with spontaneous behavioral monitoring | - | - | - | - | 11 |
| Verifying experiment 2: Fiber photometry with free-moving behavior | - | - | - | - | 6 |
| Verifying experiment 3: 30-Hz optogenetic stimulations of BF PV neurons for fMRI and behaviors | - | 12 | - | - | - |
| Comparing artifacts of PMMA and SiO<sub>2</sub> optical fiber | - | - | - | - | 5 |
| Anatomical tracing of BF ChAT neurons using ChrimsonR-mCherry | 4 | - | - | - | - |

### METHOD DETAILS

#### Stereotaxic surgeries

All stereotaxic surgeries were conducted under isoflurane anesthesia (5% for induction and 1-1.5% for maintenance), and animals were placed on a stereotaxic frame with a heating pad (mouseSTAT, Kent scientific cooperation). A midline sagittal incision was made along the scalp to expose the skull. Two craniotomies (∼0.4 mm in diameter) were made on top of bilateral BF (AP 0 mm, ML ±1.3 mm), targeting the intermediate portion of the BF, including the horizontal limb of the diagonal band of Broca (HDB), magnocellular preoptic nucleus (MCPO) and substantia innominate (SI). AAV2/9-hSyn-DIO-ChrimsonR-mCherry-WPRE-hGH pA (Titer: 4.76 × 10^12^ v.g. /mL; BrainVTA Co., China) was injected into each side of BF(AP 0 mm, ML ±1.3 mm, DV -5.0 mm from dura) utilizing Nanoject III (Drummond Scientific) via a glass pipette (1 nL/s, 20 nL/cycle, inter-cycle interval 10 s, total 20 cycles and 0.4 µL). After each injection, the needle was slowly pulled up following a 10-min waiting period to prevent backflow. Then scalp was sutured, and mouse was placed in a postoperative incubator for anesthesia recovery.

Optical fiber and head holder implantation for awake optogenetic fMRI were performed two weeks after virus injection. Head holder design and the surgical procedure were modified from our previous study^26^. The scalp over the skull was removed, and then the periosteum was removed with cotton swabs dipped in 3 % hydrogen peroxide. The tissue covering the skull posterior to the lambda point was also removed for firmer fixation of the head post. Then, the exposed skull was cleaned by saline. After drying the surface of skull, a coat of self-etch adhesive (3 M ESPE Adper Easy One) was applied followed by light curing.

To reduce imaging artifacts caused by fiber implantation and also accommodate the limited space of the cryogenic RF coil, we chose flexible fiber optic implants (200 µm in diameter; NA 0.5; Inper Co., China) made of poly-methylmethacrylate (PMMA) which has similar magnetic susceptibility to the brain tissue/water^60^. Two craniotomies (∼0.4 mm in diameter) were made on top of bilateral BF and the PMMA optic fibers were inserted into bilateral BF (AP 0 mm, ML ±1.3 mm, DV -4.8 mm from dura). After draining the overflow of liquid from the craniotomy with dust-free paper, the light curing flowable dental resin was applied and cured with blue light for 20 s to fix the optic cannula to skull. After adhering the skin surrounding the exposed skull with tissue adhesive, the dental resin was applied on the occipital bone, and a custom-made MRI-compatible head holder was place closely on the same place. Then, the flexible PMMA optic fibers were bent and fixed on the surface of skull and head holder with dental resin (Figure 1B). Finally, other exposed regions of the skull was covered with a thick smooth layer of dental cement. After surgery, the mice were placed in a postoperative incubator for anesthesia recovery.

#### Habituation for awake mouse optogenetic fMRI (opto-fMRI)

After 1-week recovery from the fiber and head holder implantation, mice were habituated for awake fMRI for seven days (Figure 1A). Mice were head-fixed on an animal bed with the recorded acoustic MRI scanning noise, following a previously described habituation paradigm ^28^. The detailed schedule was listed in below table. No reward was given during and after the habituation training.

**Table.** Summary of habituation schedules for awake mouse ofMRI.

|  | Day1 | Day2 | Day3 | Day4 | Day5 | Day6 | Day7 |
| --- | --- | --- | --- | --- | --- | --- | --- |
| **Duration of Habituation** | 60 min | 60 min | 60 min | 60 min | 60 min | 60 min | 60 min |
| **Acoustic Noise** | - | 60 dB | 60 dB | 90 dB | 90 dB | 110 dB | 110 dB |

“–” denote absence.

#### Optogenetic fMRI acquisition

All MRI data were acquired with a Bruker BioSpec scanner (9.4T/30cm, Bruker, Billerica, USA, ParaVision 6). An 86 mm volume coil was used for transmission and a 4-channel cryogenic phased array mouse head coil (Bruker) was used for receiving. A thick layer of Kwik-cast was applied smoothly over the mouse skull immediately before the MRI scanning to reduce distortion caused by air. The mouse was first secured in the animal bed without any anesthesia. Light delivery systems were kept outside the magnet and 7 m fiber optic patch cable (200 µm in diameter; NA 0.5; Doric Lens) were used to deliver light into the bore of the scanner. Optical patch cable was attached to the implanted optic fiber using a zirconia sleeve (Thorlabs). Then the mouse was placed inside the MRI scanner. A T2-weighted structural image was acquired for co-registration with following parameters: TRL=L3400 ms, TEL=L33 ms, field of view = 16 × 16 mm^2^, matrix size = 256 × 256, slice thickness = 0.5 mm, 30 axial slices, RARE factor = 8, and number of averages = 2. After field map based local shimming within the mouse brain, Biband-EPI was acquired using the multiband EPI sequence ^28^ with the following parameters: TR = 500 ms, TE = 15 ms, flip angle = 38.8°, bandwidth = 300 kHz, field of view = 15 × 12 mm^2^, matrix size = 100 × 80, nominal slice thickness = 0.48 mm (slice thickness 0.4 mm with a gap of 0.08 mm), 898 or 982 volumes (scan 1-3: 898 volumes and scan 4-6: 982 volumes), and 38 axial slices per scan. During the Biband-EPI scanning, an Arduino Uno board (https://www.arduino.cc/) was use to synchronize the scanner trigger and the lasers for optogenetic stimulations. Red light (10 Hz, 5% duty cycle) was delivered to the bilateral BF utilizing 638 nm laser (Shanghai Laser & Optics Century Co., Ltd.), and the power at the fiber tip was 5 mW. The overall stimulation setup was shown in Figure 1D. For each EPI run, 28 stimuli were delivered with durations of 0.5 s or 2 s, and a random inter-trial-interval time of 15 +/- 5 s.

#### Simultaneous behavioral monitoring and fMRI acquisition

For the awake mouse fMRI with simultaneous behavior monitoring, a similar imaging protocol was used as mentioned above, with the following difference: Biband-EPI, TR = 750 ms, flip angle = 46.6° and 10000 volumes per scan. In addition, two custom-made MR-compatible video cameras (sampling rate of 25 fps, 1080 × 980 pixels) was placed inside the bore to record the spontaneous behaviors, e.g., grooming and body motion, which were later used to estimate the grooming evoked BOLD activations.

#### fMRI data processing

After the image format conversion, the mouse brain was extracted manually utilizing ITK-SNAP (http://www.itksnap.org/). All subsequent procedures were performed utilizing custom scripts in MATLAB 2020a (MathWorks, Natick, MA) and SPM12 (http://www.fil.ion.ucl.ac.uk/spm/). Images of each scan were registered to the scan-specific structural image utilizing rigid body transformation, and the scan-specific structure image was then nonlinearly transformed to the 3D Allen Mouse Brain Common Coordinate Framework, v3 (CCFv3, http://atlas.brain-map.org/) for group analysis. Then, a light spatial smoothing (0.4 mm isotropic Gaussian kernel) was performed. Furthermore, BOLD signals were regressed by “6 rp + 6 Δrp + 10 PCs” nuisance signals to minimize the effects of scanner drift, motion and other non-neural physiological noises (Figure S2), adopted from our previous study ^26^. “6 rp + 6 Δrp” nuisance signals represented 6 head motion parameters and their 1^st^ order first derivatives, and “10 PCs” were the first 10 principal components from the BOLD signals of non-brain tissue, e.g., the muscles. The Pearson’s correlation coefficients between the frame-wise displacement (FD) and DVARS (D referring to temporal derivative of time courses, VARS referring to RMS variance over voxels) were calculated to quantitatively evaluate to what extent the motion related signal was reduced by given regressors ^29^.

#### General linear model of cell-type specific optogenetic stimulations and spontaneous behaviors

GLM based statistical analysis was conducted utilizing the awake mouse-specific hemodynamic response function (HRF) from our previous study^26^. Standard first level analysis was done for individual EPI scans. For the awake mouse fMRI experiment with optogenetic stimulations, the period of optogenetic stimulations was set as the predictors. For the second level analysis, we used a flexible factorial model including cell-type specific optogenetic stimulations, i.e., VGLUT2 (or ChAT, PV, SOM) v.s. Control, and individual mouse (as random effect). The resulting activation maps were thresholded with FDR corrected p < 0.05.

For the awake mouse fMRI experiment with simultaneous behavior monitoring, the behaviors (e.g., grooming and body motion) were used as the predictors and thus the remaining period was used implicitly as the baseline. A one-sample t-test was conducted for second level analysis to generate the activation maps with FDR corrected p < 0.05.

#### Specificity of DMN-like activations driven by BF neurons

The ICA-based functional networks were adopted from a previous study^77^, in which they identified macro-communities to spatially encompass six macroscopic networks of the mouse cerebrum. Briefly, the mouse resting state (rs-)fMRI dataset was fed into MELODIC (Multivariate Exploratory Linear Optimized Decomposition of Independent Components) ^78^ to perform within-subject spatial ICA. Using preset 30 components and the ICA-FIX pre-processing pipeline, 23 out of 30 components were detected in meaningful anatomical regions of the mouse brain and considered as rs-fMRI functional networks. Then, hierarchical network analysis was performed based on total correlation matrices of the 23 group ICA signal components and subsequently merged the group ICA components into six cerebral clusters, i.e., DMN-like (Figure 2B), somatosensory, sensory, hippocampus, basal ganglia and olfactory networks (Figure S7A). The proportion of significantly activated voxels that overlapped with ICA-based functional networks (Figure 2B and Figure S7A) was calculated with opto-fMRI responses were thresholded at FDR-corrected p < 0.05, while ICA-based functional networks were thresholded at z > 3.

Receiver operating characteristic (ROC) curves were computed to evaluate the specificity of opto-fMRI activations (Figure 1E and Figure 2A) on mouse ICA-based functional networks (Figure 2B and Figure S7A)^79^. Briefly, functional networks and opto-fMRI responses were normalized between 0 and 1, and were binarized using a series of 100 thresholds to keep the 0-100% of activations, respectively. The resulting binary functional networks and opto-fMRI responses were compared using the ROC analysis with the corresponding functional network, i.e., DMN-like network, as the ground truth. The true positive and false positive rate vectors were plotted against each other and the resulting area under the curve (AUC) was used as the measurement of the specificity of opto-fMRI activations on mouse functional networks. The AUC results were then compared against a null distribution of the same datasets using the permutation test with 1000 times shuffling of the cell-type specific opto-fMRI responses.

#### Anterograde axon tracing from BF neurons

The whole-brain axonal projections data from the four BF cell types is modified from our previous study^4^. Briefly, ChAT-Cre, PV-Cre, SOM-Cre or VGLUT2-Cre mice were anesthetized with ∼1.5% isoflurane in oxygen, 300-400 nL output tracing vector (AAV2-EF1a-FLEX-mCherry, titer: ∼10^12^ gc/mL) was inject into the right BF. Two to three weeks after AAV injection, mice were deeply anesthetized with isoflurane and immediately perfused intracardially with PBS followed by 4% PFA. Brain was removed and post-fixed in 4% PFA in PBS at 4 LJC overnight, dehydrated in 30% sucrose in PBS for 48 hr, embedded in Tissue Freezing Medium, cut in 30 or 50 μm coronal sections and mounted with VECTASHIELD antifade mounting medium with DAPI. All sections were imaged utilizing 20X/0.75 objective in a high-throughput slide scanner for further 3D reconstruction and quantification. The digitized analysis was performed with a software package which consists of three modules: image registration, signal detection, and quantification.

#### Voxel-wise whole-brain mesoscopic structural connectivity

Whole-brain structural connectivity was obtained from the Allen Mouse Brain Connectivity Atlas (http://connectivity.brain-map.org/), which was based on 428 viral microinjection experiments in C57BL/6J male mice. Briefly, adeno-associated viral anterograde tracers, containing genes encoding for enhanced green fluorescent protein, were sterotactically injected at different sites in mice. After the injection, 2 weeks were allowed for the protein expression before the animals were killed, the brain was extracted, sectioned, imaged with two-photon microscope, reconstructed into 3D fluorescence maps, and transformed into a common reference space ^80^. Then, the connectome data were aggregated according to a voxel-wise interpolation model ^31^, modeling the connectivity at each voxel as a radial basis kernel-weighted average of the projection patterns of nearby injections. Thus, a high-resolution mouse brain connectome (100 μm^3^) was estimated according to a Voronoi diagram based on Euclidean distance between neighboring voxels ^31^. The Voronoi-based resampling allowed us to spatially weight voxels with respect to neighboring areas, and preserve the intrinsic architectural foundation of the connectome. Finally, fiber tracts and ventricular spaces were filtered out, yielding a final weighted and directed 15,314 × 15,314 matrix composed of 0.027 mm^3^ aggregate Voronoi voxels. Notably, the structural connectivity matrix was transformed by the logarithm operation similar to our previous study ^71^.

#### Non-negative matrix factorization

To further investigate how the BF modulated global organizations of mouse cerebral functional networks, we firstly concatenated the four cell-type secondary projections from BF neurons (Figure S10A-C), in which the secondary projections was adopted from source-weighted whole-brain mesoscopic structural connectivity by cell-type output from BF neurons, thus resulting a 15,314 × 61,256 connectivity matrix (Figure S10A-C). Considering the bilateral optogenetic stimulations on BF neurons, we averaged the connectivity strengths from ipsilateral and contralateral hemispheres for both cell-type specific output from BF (Figure 4B) and whole-brain mesoscopic structural connectivity matrix. Then, the non-negative matrix factorization (NMF) was performed on the resulting secondary connectivity matrix (15,314 × 61,256, Figure 4B). The spatial NMF components (*W_p_*_0×15,314_) were back-projected to whole-brain BOLD signals to reconstruct the time course of each component (*S_t_*_×*p*0_) for each individual EPI scan. NMF was chosen over other dimensional reduction methods, e.g., PCA or ICA, because it enhances the interpretability of the resulting low dimensional components. To estimate the appropriate dimensionality of the NMF analysis, we calculated the mean cophenetic coefficients following the NMF decomposition (a measure of the optimal number of subgroups in the connectivity matrix), and found the maximum coefficient in the top 6 components (Figure S10D). Thus, we chose the top 6 NMF components in all following analysis.

#### Optogenetic free-moving behavioral test

Free-moving optogenetic behavioral test was performed in a 40 × 40 × 40 cm^3^ arena (Figure 5A). To keep the behavioral testing condition similar to the MRI experiment, the phase of familiarization was set to 30 min with two familiar objects in two corners of arena. Then, one of the familiar objects was replaced by a novel one (different shape, texture, size, and color). The same optogenetic stimulations as in MRI were then delivered to the bilateral BF immediately (Figure 5B). Animal behaviors were recorded utilizing a video camera (25 fps, 1080 × 980 pixels), which was positioned 50 cm above the center of the arena floor.

To analyze the recorded video, a set of behaviors including (1) locomotion speed, (2) quiet wakefulness, (3) grooming, (4-5) familiar and novel object explorations were recorded. Object exploration was defined as the mouse’s nose was < 1 cm from the object and the dwell time was < 4 s. Locomotion was labelled when the speed was > 2 cm/s excluded the period of object explorations. Moreover, we fitted the outline of mouse body by an ellipse and used the major (*a*) and minor (*b*) axes of the ellipse to distinguish the grooming and quiet wakefulness. The behavioral state of grooming was termed when the mouse was in a quasi-stationary condition (speed < 2 cm/s) and the outline of mouse body was sub-rounded (i.e., *a* – *b*> 2 cm). The remaining quasi-stationary behavior (speed < 2 cm/s, *a* – *b* < 2 cm) was defined as the quiet wakefulness with short periods (dwell time < 5 s) excluded. Durations of the five types of behaviors were scored.

#### Decoding the global patterns of mouse behaviors

A general decoding model was used to predict the cerebral responses of mouse behaviors (Figure 6B & S16). As the optogenetic fMRI and free-moving behavioral tests were conducted in different days, we divided the total behavior length into 6 runs for each mouse, keeping temporal alignment with the order of fMRI scanning. And also, the BOLD activation maps were decomposed by the NMF strategy. Then, we averaged both the weights of NMF components and behavioral performances across individual mice, and only considered the group-wise behaviors under optogenetic stimulations on cell-type specific neurons. The resulting cell-type specific BOLD weights of NMF components (*W*_5×6×6_) were set as the model predictors (5 groups, 6 runs and 6 NMF components), and the behavioral performances (*Y*_5×6_) were termed as the model output.

We randomly divided 6 runs into two groups to generate encode and decode datasets respectively, and repeated 20 times for traversing all random conditions, i.e., choose 3 runs as encoding dataset from all 6 runs. The regularization penalty was estimated on the encoding dataset utilizing marginal maximum likelihood estimation with minor modifications that reduced numerical instability for large regularization parameters. To avoid over fitting, ridge regression was chosen over other regularization methods (e.g., LASSO or elastic net) because a sparseness prior would isolate the best contributing model variables and completely reject variables which have slightly less informative. Regression coefficients (*β*_0_, *β*_1_, *β*_2_, *β*_3_, *β*_4_, *β*_5_ and *β*_6_) were termed as the representative vector [*β*_1_, *β*_2_, *β*_3_, *β*_4_, *β*_5_, *β*_6_] for each mouse behavior in NMF space, and further used to construct corresponding spatial patterns ([*β*_1_, *β*_2_, *β*_3_, *β*_4_, *β*_5_, *β*_6_]·*W*_6×15314_) by multiplying the spatial NMF components (*W_p_*_0×15,314_ and *p*_0_ = 6).

#### Histology and microscopy

To verify the expression of the virus and placements of the optical fibers, mice were deeply anesthetized and immediately perfused utilizing 0.1M phosphate-buffered saline (PBS), followed by 4% paraformaldehyde (PFA) in PBS. The brain tissues were removed and post-fixed overnight in 4% PFA. Subsequently, the brains were transferred to a 30% sucrose solution at 4 °C for 48 hr. Brain samples were embedded with OCT compound (NEG-50, Thermo Scientific) and cut into 50-μm sections utilizing a cryostat (HM525 NX, Thermo Scientific). Brain sections were stored at -20 °C before further processing. For histology examination, brain sections were washed in PBS and coverslipped with mounting media containing DAPI (# F6057, Sigma). The fluorescence images were captured utilizing an epifluorescence microscope (VS120, Olympus).

#### Free-moving behavioral tests with simultaneous fiber photometry

To verify the predicted behavior-related cerebral patterns, the neuronal activity of primary motor area (MOP), entorhinal area (ENT) and anterior cingulate area (ACA) were recorded using calcium fiber photometry during spontaneous behavior in the same arena as above. AAV2/9-hSyn-GCaMP6s-WPRE-hGH pA (titer: 2.69×10^12^ v .g. /mL; BrainVTA) was injected into unilateral MOP (AP +0.98 mm, ML +1.5 mm, DV -0.54 mm from dura), ENT (AP -3.88 mm, ML +3.75 mm, DV -2.3 mm from dura) and ACA (AP +0.14 mm, ML -0.3 mm, DV -1.07 mm from dura). Three optic fibers (200 µm in diameter; NA 0.48; Newdoon Co., China) were inserted into the three regions respectively, and secured with the light curing flowable dental resin. Finally, the exposed skull was covered with dental cement.

Four weeks after the surgery, the same free-moving behavioral test as above was performed with simultaneous fiber photometry recording and behavioral monitoring (Figure 7E). Three patch cables were connected to the three fiber implants with sleeves. Three 465 nm LEDs (CLED_465, Doric Lens) were used as excitation sources. Excitation power at the tip of the fiber was ∼20 μW. Three fluorescence optical mini cubes (FMC5_E1(465-480)_F1(500-540), Doric Lens) and three visible femtowatt photoreceivers (NPM_2151_FOA_FC, Doric Lens) were used for the fluorescence detection. During recording, a software-controlled lock-in detection algorithm was implemented in the TDT RZ2 system utilizing the fiber photometry ‘Gizmo’ of the Synapse software (https://www.tdt.com/). The photometry signals were recorded at a sampling rate of 1017 Hz.

To analyze the photometry data, a median filter with 10-min window length was applied to the raw GCaMP6s photometry signals to capture the infra-slow baseline drift. The median-filtered calcium signal was then regressed out from the raw photometry signals using the least-squares regression. Then, the changes of calcium signals were quantified as (F-F0)/F0, where F was fluorescence intensity at each time point and F0 was calculated by mean minus double standard deviations. The calcium time series were further down-sampled to 25 Hz for subsequent analysis using the MATLAB function *interp1()*. Finally, the neuronal firing rate was evaluated by deconvolution of the normalized calcium signals (F-F0)/F0 using the first-order auto-regressive model.

For behavioral state classification, the same criteria was applied to label the spontaneous behavioral state of each mouse (details in the section of Optogenetic free-moving behavioral test).

#### Optogenetics with simultaneous fiber photometry

To further explore why the locomotion and BOLD activations appeared incoherent in sensorimotor regions, we conducted the calcium fiber photometry to record opto-evoked neuronal calcium activities in SSp-m (the negatively activated region) and RSP (the positively activated region) for VGLUT2 and Control group. Bilateral BF was injected with AAV2/9-hSyn-DIO-ChrimsonR-mCheery-WPRE-pA, and unilateral SSp-m (AP +1.0 mm, ML +2.75 mm, DV -0.9 mm) and RSP (AP -3.8 mm, ML +1 mm, DV -0.5 mm) were injected into AAV9-hSyn-GCaMP6m-WPRE-hGH-pA (titer: 2.69×10^12^ v .g. /mL; Shanghai Taitool Bioscience). The same optogenetic stimulation conditions used in MRI experiments were applied to both sides of the BF. The recording and analysis of calcium data followed the aforementioned protocol.

#### Significance test using the spatial autocorrelation preserving shuffling

Because profiles of predicted maps of mouse behaviors (Figure 6G) are spatially auto-correlated, we adopted a procedure from previous studies to overcome this issue and generate statistical significance^81–83^. Briefly, we generated surrogate maps that randomly varied in their particular topographies (nL=L10000 times shuffling) but preserved the general spatial autocorrelation structure (Figure S17B-C). Using null distributions generated from spatial autocorrelation preserving surrogate maps, we determined the significance level, i.e., *p*_spin_, of spatial correlation between empirical grooming map and the predicted maps of mouse behaviors (Figure 7D & S17A).

#### Comparing imaging artifacts of PMMA and SiO_2_ fiber optic implants

A separate cohort of wild-type mice was utilized to compare imaging artifacts induced by PMMA and SiO_2_ fiber optic implants (200 µm in diameter; NA 0.48; Inper) inserted unilaterally into the basal forebrain. To accommodate the limited space of the cryogenic RF coil, the ceramic ferrules of SiO_2_ fiber optic implants were trimmed. All other surgical procedures and fMRI acquisition parameters were identical to those described above. The artifact size caused by the fiber optic implant in T2 or EPI images was measured, and a two-tailed paired t-test was employed to determine statistical significance.

### QUANTIFICATION AND STATISTICAL ANALYSIS

Statistical analyses were performed based on Matlab (Mathworks, Natick, MA). Statistical significance was determined by two-tailed or paired Student’s t test for comparisons between two groups and by ANOVA with Tukey-Kramer’s test for comparisons among multiple groups. Correlation between groups was tested by Pearson’s correlation. Statistical details, including p values and sample numbers, were described in the relevant methods sections or figure legends.

