## Supplemental Figures and Tables for "Cell-type specific optogenetic fMRI on basal forebrain reveals functional network basis of behavioral preference"

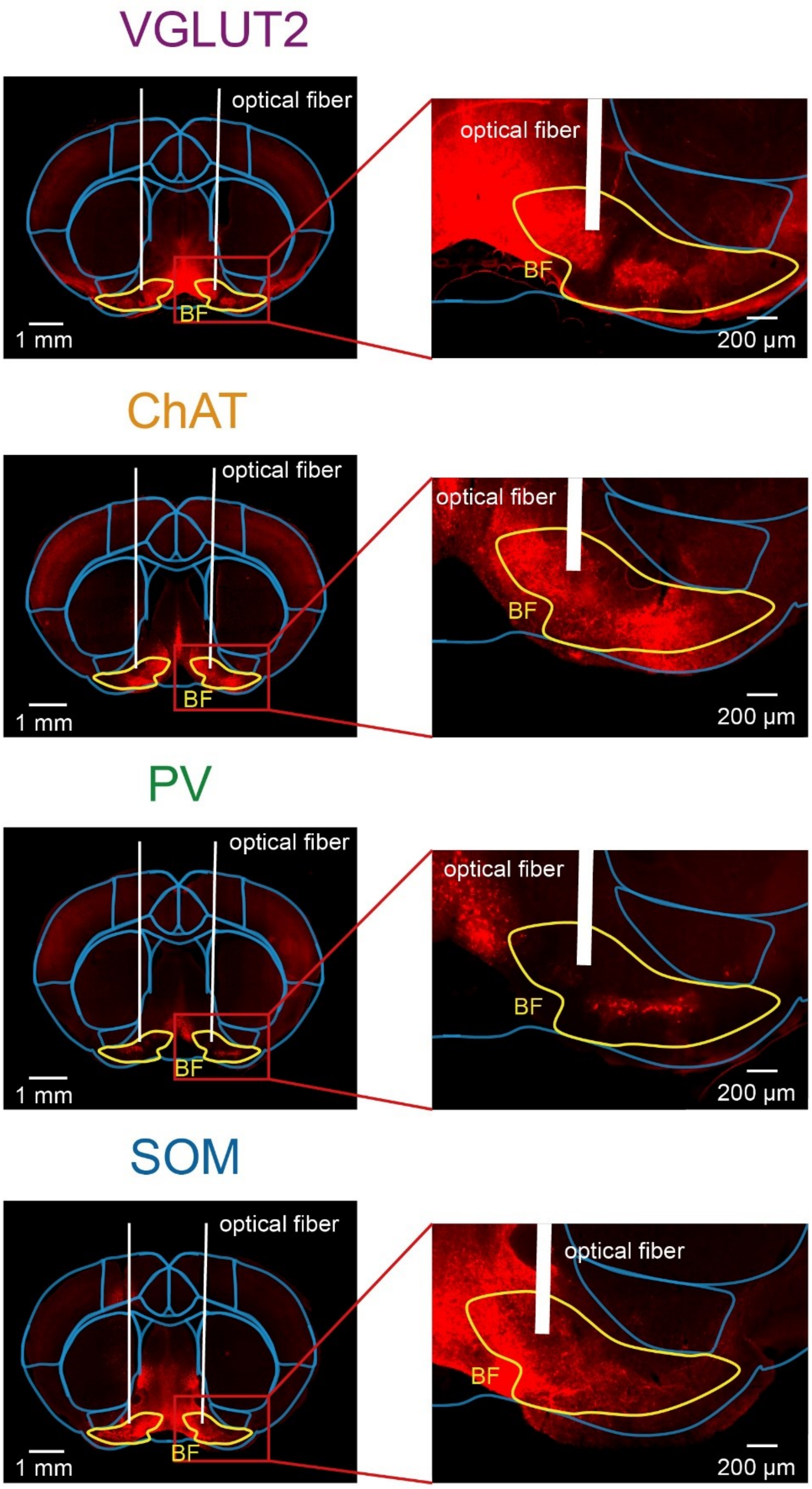


**Supplementary Figure 1 | Related to Main Figure 1**

**Histological verification of virus expressions in basal forebrain (BF) neurons.**

VGLUT2, ChAT, PV and SOM refer to four cre-line mice that are specific to different cell types of BF. VGLUT2, vesicular glutamate transporter 2; ChAT, choline acetyltransferase; PV, parvalbumin; SOM, somatostatin.

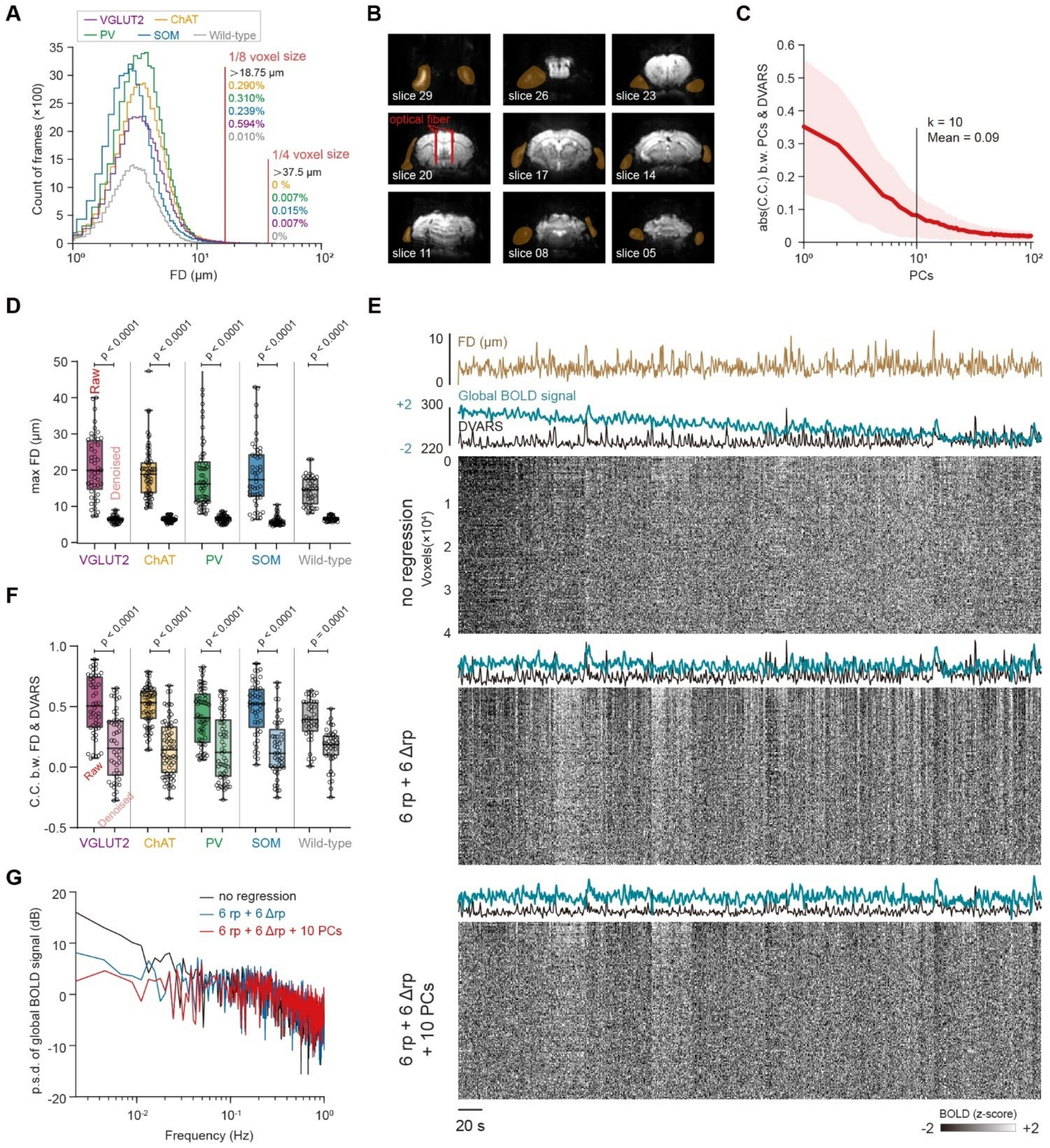


**Supplementary Figure 2 | Related to Main Figure 1**

**Regression of “6 rp + 6 rp +10 PCs” notably reduced the impact of nuisance signals on whole brain BOLD signals.**

(A) Small head motion level of opto-fMRI sessions during awake mouse fMRI scanning. Notably, the in-plane spatial resolution of EPI images was 150 μm. FD, frame-wise displacement.

(B) Representative mask (orange shadows) used for extracting the principal components (PCs) outside mouse brain tissue to model the non-neural nuisance signals.

(C) First 10 PCs highly captured the nuisance signals during the fMRI scanning. Higher correlation between PCs and DVARS (the temporal Derivative of root mean square VARiance over voxelS) represented more head motion effects on BOLD signals were captured by PCs.

C.C., Pearson's correlation coefficients. Red line (or shadow), mean (+/- SEM.) correlation.
(D) Significant decrease of head motion level after denoising approach based on the "6 rp + 6 rp +10 PCs" regression. Statistical significance was calculated by two-tailed paired t-test. Each dot represented an individual EPI run.
(E) Visualization of effects of nuisance signal regression. Representative data with head motion and resulting signal variations. The time series of whole brain voxels was shown as intensity plots. Each dot represented an individual EPI run.
(F-G) Minimal head motion effects (F) and infra-slow drift (G) of BOLD signals after the "6 rp + 6 rp +10 PCs" regression. Statistical significance was calculated by two-tailed paired t-test. p.s.d., power spectral density.

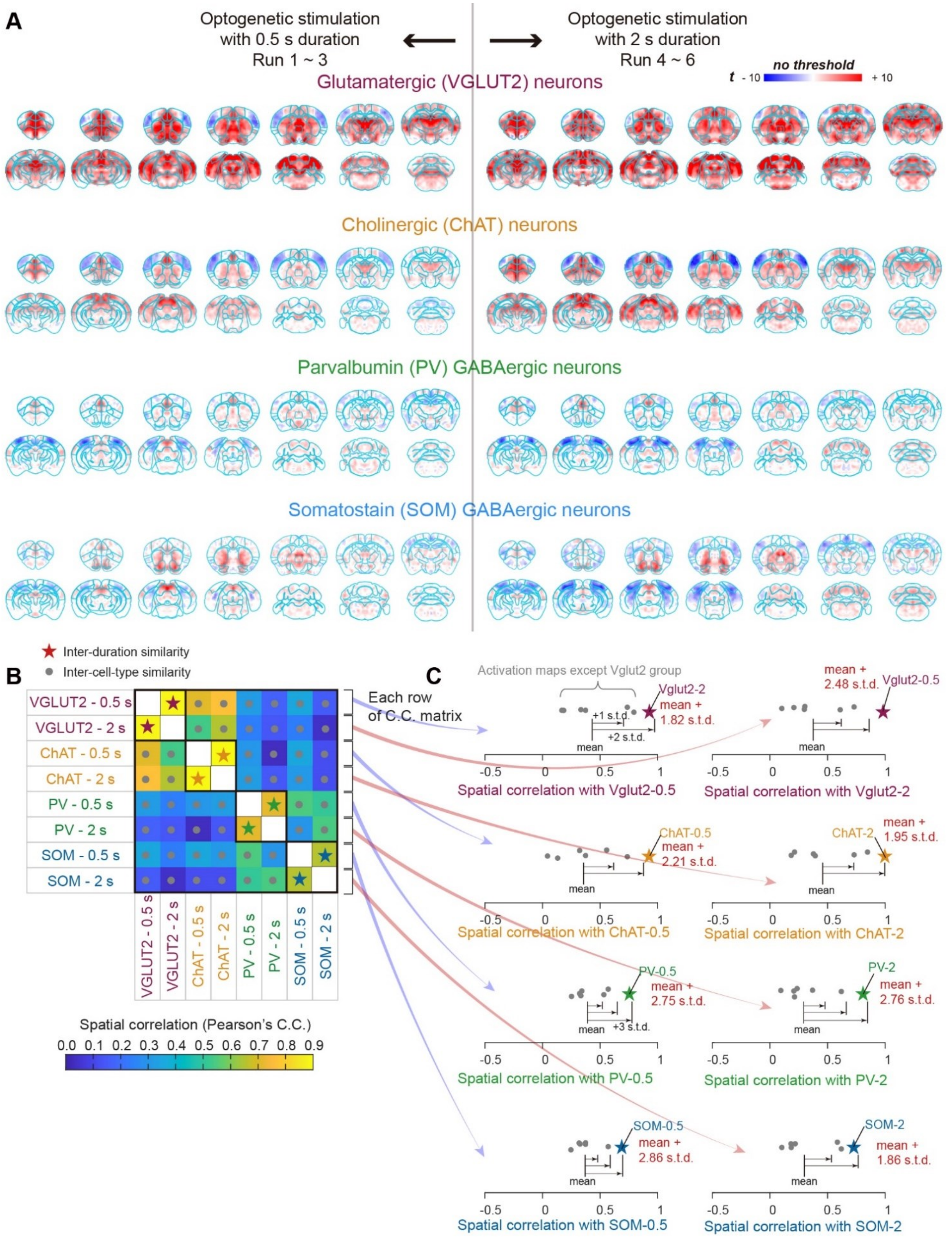


*(Legend on next page)*

**Supplementary Figure 3 | Related to Main Figure 1**

**Spatial similarities of inter-duration BOLD activation maps were significantly higher than those of inter-cell-type ones.**

(A) Functional activation maps under optogenetic stimulations of cell-type specific BF neurons. BOLD activation maps were generated using the two-sample t-test, i.e., VGLUT2 (or ChAT or PV or SOM) v.s. Control, respectively. Activation maps were showed without thresholds for both 0.5-s duration (left) and 2-s duration (right) optogenetic stimulations.

(B) High spatial similarities of the activation maps between 0.5-s duration and 2-s duration for four BF cell-types. The color of each square indicates the spatial similarity (Pearson's correlation coefficients, C.C.) among opto-fMRI activation maps.

(C) Spatial similarities of inter-duration BOLD activation maps were significantly higher than those of inter-cell-type ones. Notably, for each BOLD activation map, the inter-duration spatial correlation (represented by colored pentangle) was higher than all other inter-cell-type spatial correlations (represented by gray dots).

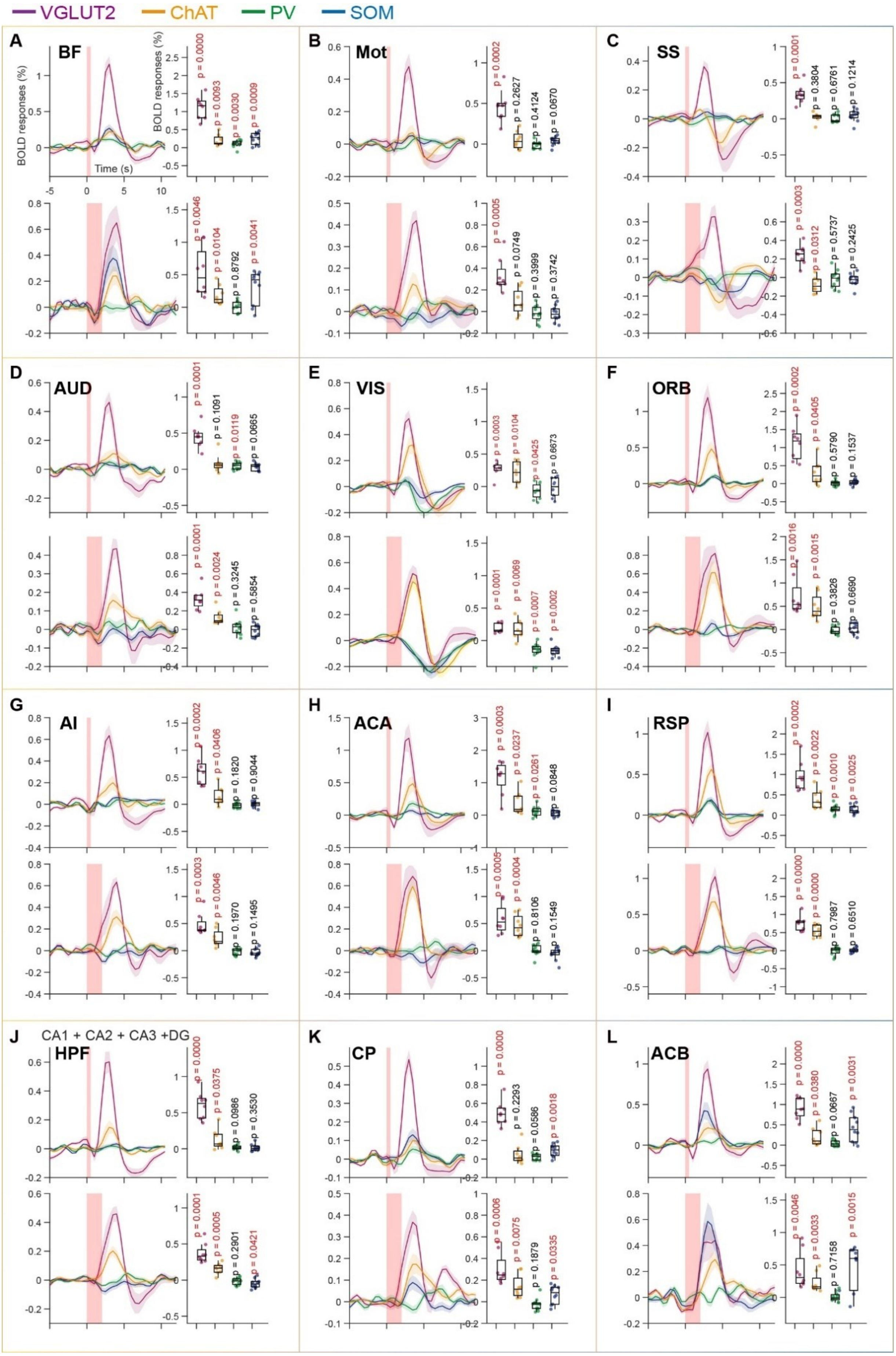


*(Legend on next page)*

**Supplementary Figure 4 | Related to Main Figure 1**

**Averaged BOLD responses drove by cell-type specific BF activations in awake mice.**

(A-L) BOLD responses extracted from regions-of-interest (ROIs) defined in Main Figure 1F. Abbreviation of each ROI was listed in Supplementary Table1.

(Left panel) Averaged BOLD signals under 0.5 s (upper panel) and 2 s (lower panel) optogenetic stimulations on BF neurons. Red rectangular shades indicated the stimulation periods. Colored lines (or shadows), mean (+/- SEM) BOLD response.

(Right panel) Quantitative measurement of BOLD responses drove by cell-type specific BF activations. Statistical $p < 0.05$ represented significant BOLD activations in the corresponding region. Each dot represented an individual EPI run. Statistical significance was calculated by two-tailed one sample t-test.

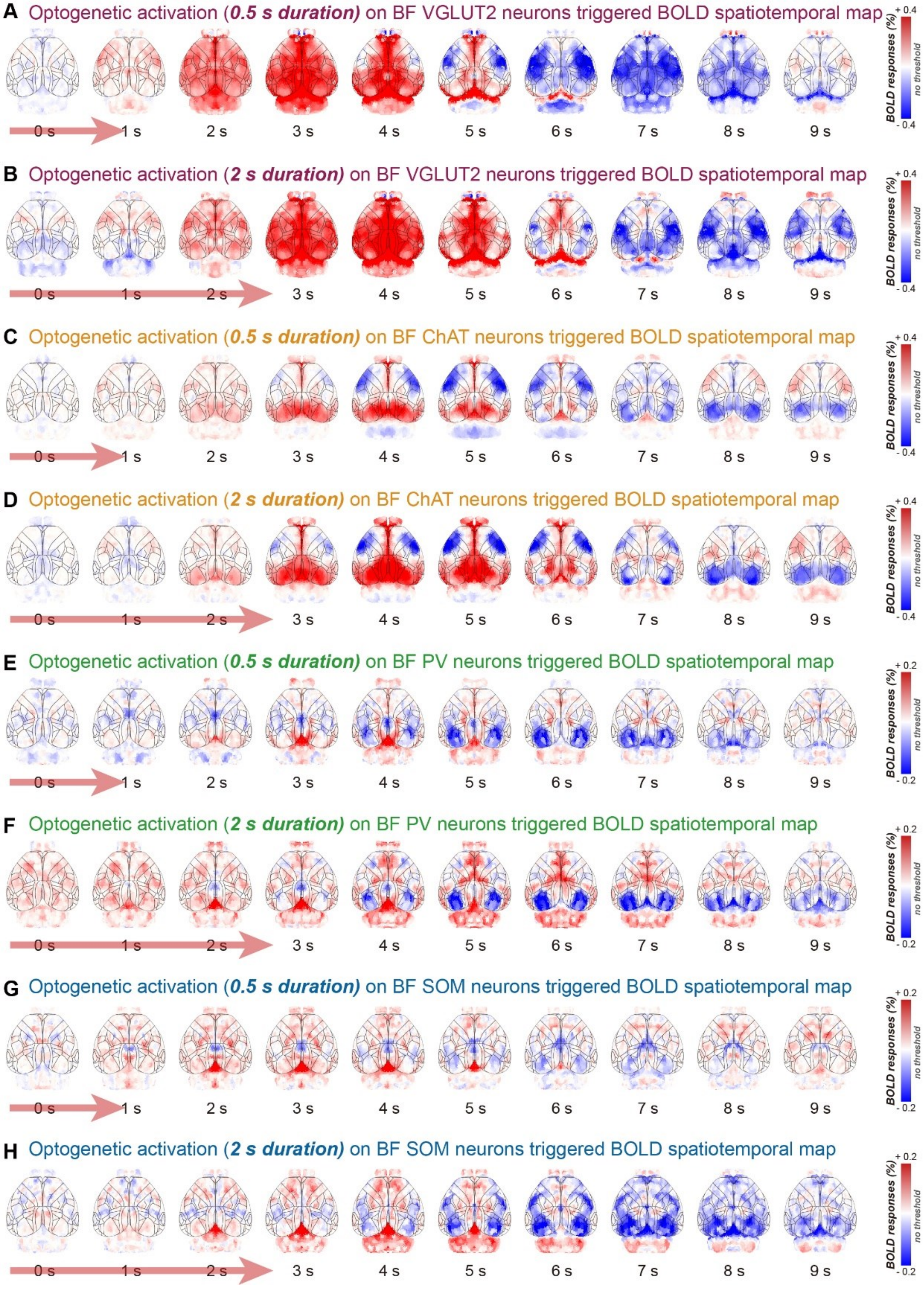


*(Legend on next page)*

**Supplementary Figure 5 | Related to Main Figure 1**

**Optogenetic activation of BF neurons evoked whole brain BOLD dynamics.**

Averaged cell-type specific opto-fMRI responses displayed on the 3D surface with the parcels based on CCFv3 Allen mouse brain atlas (black lines). Red arrows represented the 0.5-s (A, C, E and G) or 2-s (B, D, F and H) optogenetic stimulations. Notably, the opto-fMRI responses for four cell-types were subtracted by those for control group, respectively.

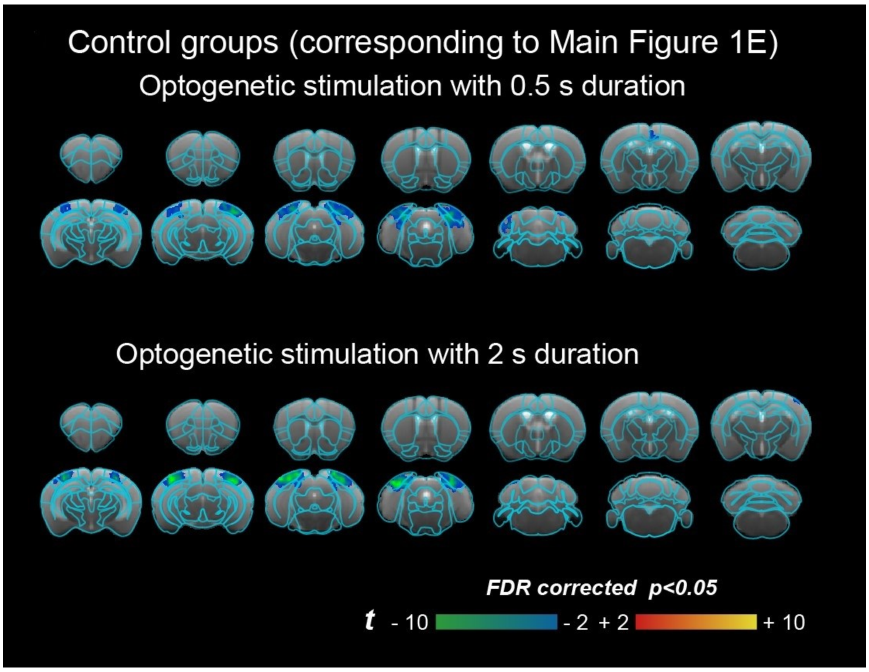


**Supplementary Figure 6 | Related to Main Figure 1**

**BOLD responses under optogenetic stimulations of BF neurons in Control group.** BOLD activation maps were generated using the one-sample t-test with FDR corrected $p < 0.05$ threshold for both 0.5-s duration (upper) and 2-s duration (lower) optogenetic stimulations.

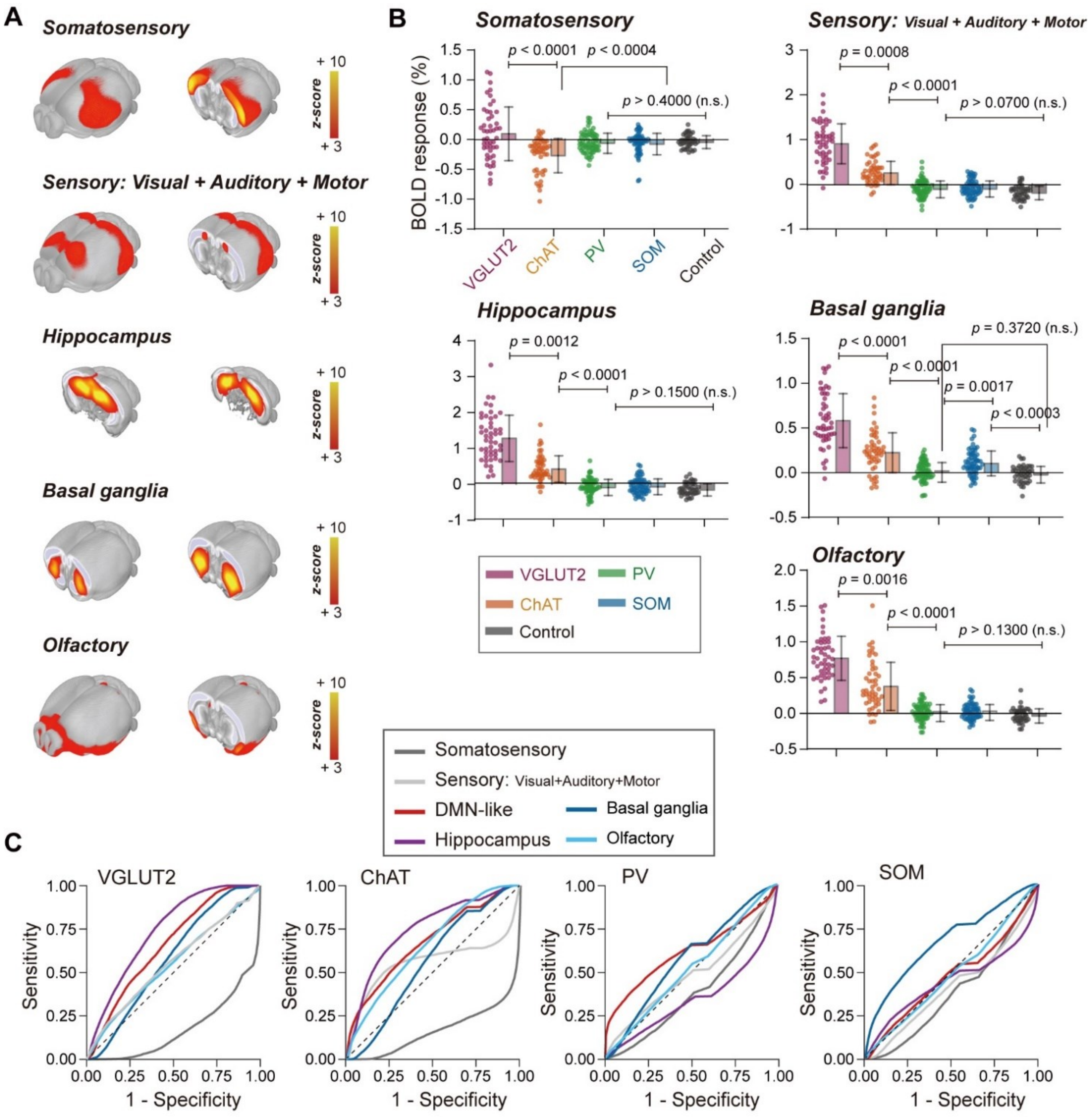


**D**

| | VGLUT2 | ChAT | PV | SOM |
|---|---|---|---|---|
| Somatosensory | 0.17 (*n.s.*) | 0.17 (*n.s.*) | 0.39 (*n.s.*) | 0.39 (*n.s.*) |
| Sensory (Visual + Auditory + Motor) | 0.55 (*n.s.*) | 0.55 (*n.s.*) | 0.45 (*n.s.*) | 0.42 (*n.s.*) |
| Default mode-like | 0.65 ($p < 0.0001$) | 0.67 ($p < 0.0001$) | ★ 0.61 ($p < 0.0001$) | 0.48 (*n.s.*) |
| Hippocampus | ★ 0.72 ($p < 0.0001$) | ★ 0.73 ($p < 0.0001$) | 0.36 (*n.s.*) | 0.43 (*n.s.*) |
| Basal ganglia | 0.58 ($p < 0.0001$) | 0.56 ($p < 0.0001$) | 0.57 ($p < 0.0001$) | ★ 0.67 ($p < 0.0001$) |
| Olfactory | 0.54 ($p < 0.0001$) | 0.65 ($p < 0.0001$) | 0.52 ($p < 0.0001$) | 0.49 (*n.s.*) |

*(Legend on next page)*

**Supplementary Figure 7 | Related to Main Figure 2**

**VGLUT2 neurons in BF evoked highest BOLD responses but PV neurons in BF modulated highest specificity of DMN-like activations in awake mouse.**

(A) Additional components in the 3D view, which were identified with independent component analysis (ICA) on resting state fMRI data. The clustered functional networks were modified from Zerbi et al.'s work in 2015[1].

(B) Highest BOLD activations across mouse functional networks drove by glutaminergic (VGLUT2) BF neurons compared to other cell-types of BF neurons. Each dot represented an individual session. Error bars, standard deviations (S.D.) of the mean. Statistical significance was determined using one-way ANOVA followed by Tukey's post hoc test for group comparison. Error bar, the standard error of the mean.

(C) Opto-fMRI activation patterns compared with mouse ICA-based functional networks[1] with ROC analysis.

(D) The specificity of opto-fMRI activations characterized by the area under the curve (AUC). A permutation test was performed by shuffling cell-type specific opto-fMRI responses 1000 times to generate a null distribution.

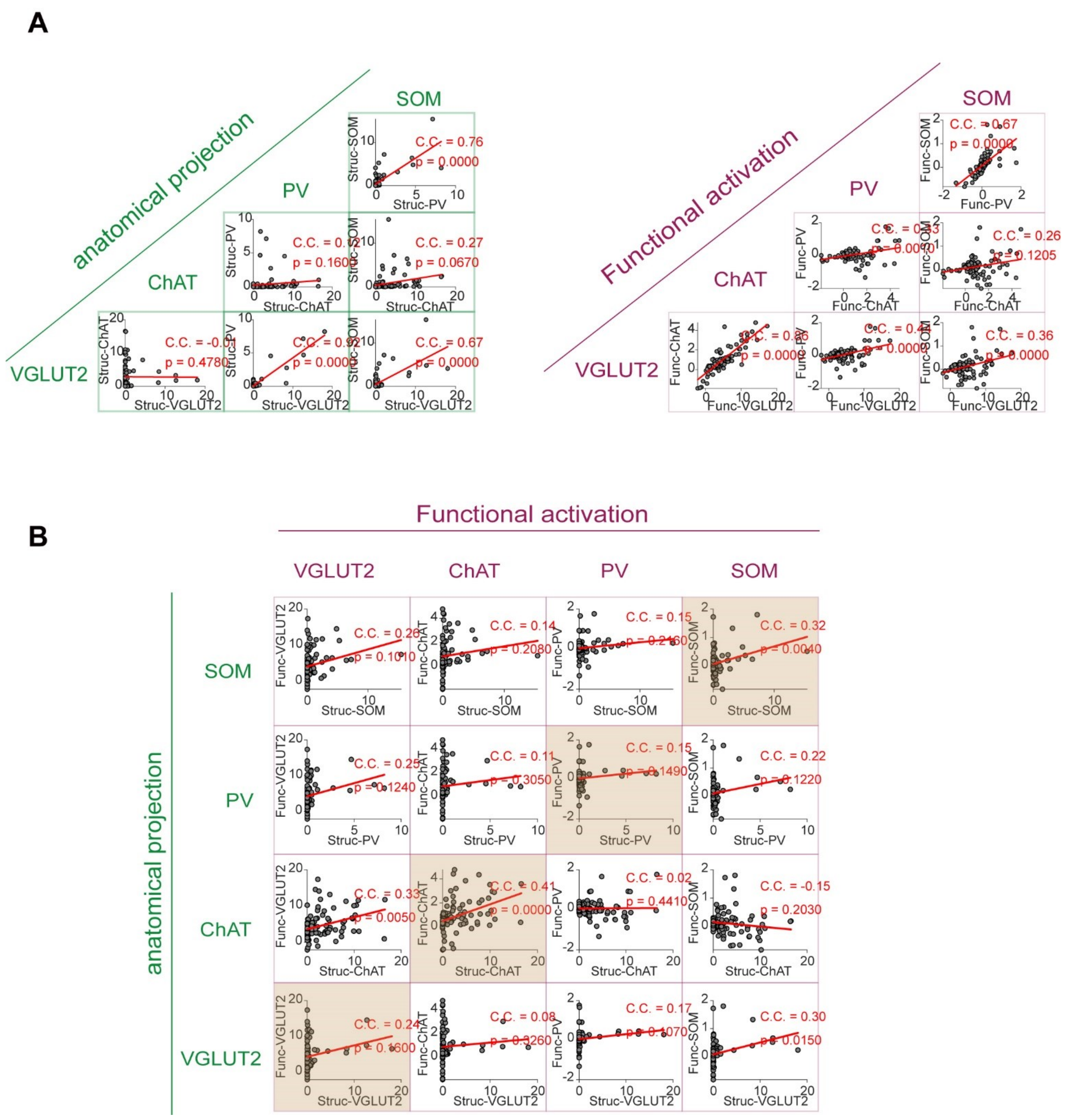


**Supplementary Figure 8 | Related to Main Figure 3**

**Spatial similarity among BF originated anatomical projections and BOLD activation maps.**

(A) High spatial similarity among the cell-type specific anatomical projections (left panel) or BOLD activation maps (right panel) of BF neurons.

(B) Weak spatial similarities (brown shadows) between anatomical projections and functional activation maps for four BF neurons.

Each dot represented an individual brain region (N = 213). The parcellation of 213 anatomical brain regions was adopted from the mesoscale mouse connectome[2]. C.C., Pearson's correlation coefficients. Red line represented the best linear fit. Statistical significance was calculated by two-tailed one sample t-test.

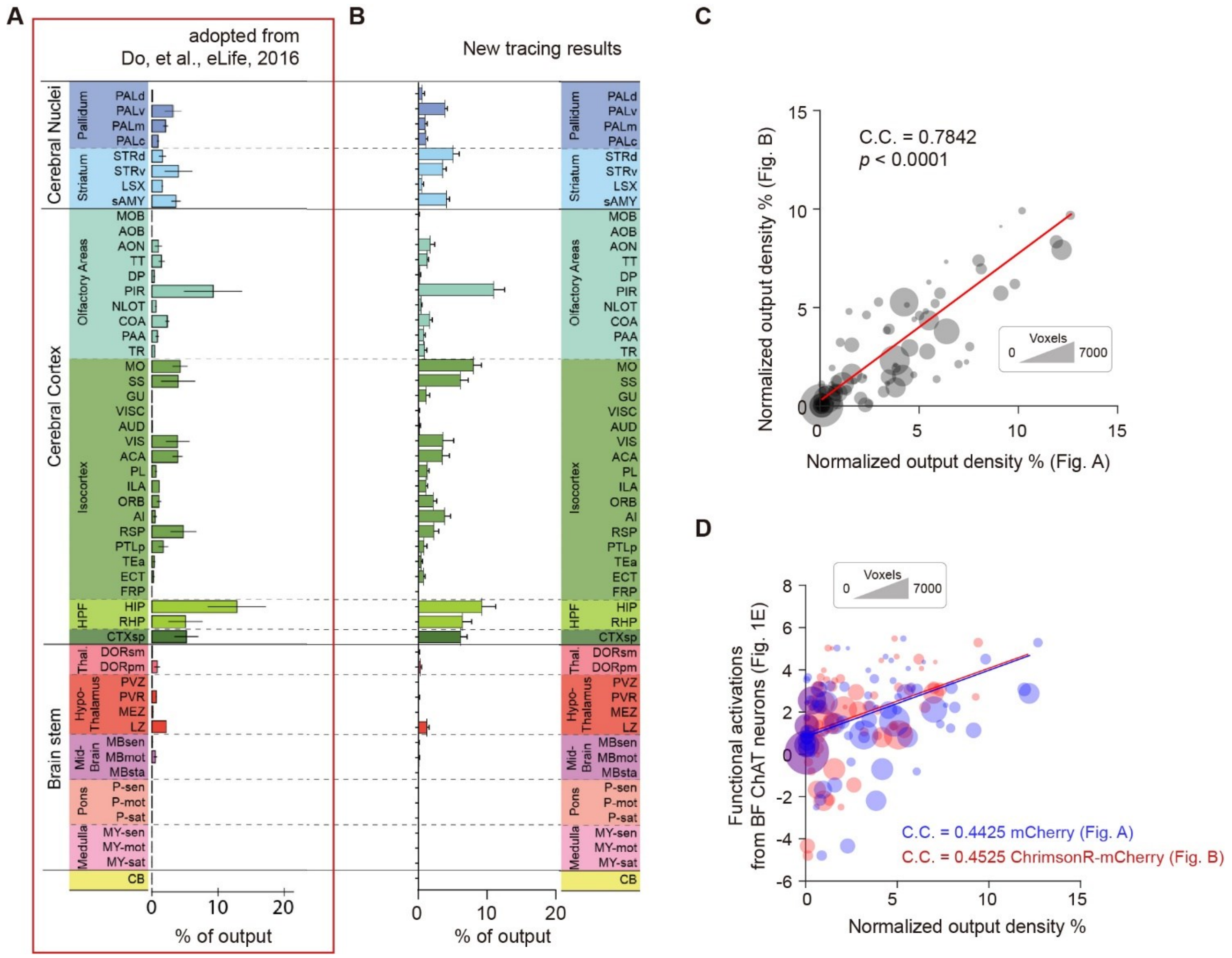


**Supplementary Figure 9 | Related to Main Figure 3**

**No significant difference in terms of axon distribution between optogenetic protein and fluorescence protein expressed ChAT-Cre mice.**

(A) Same as Figure 3B, i.e., whole-brain anatomical distribution of BF ChAT neurons based on the virus: AAV2-EF1a-FLEX-mCherry. Error bar, ± s.e.m. The bar graph was adopted from Do, et al., eLife 2016[3].

(B) Similar to (A), but based on the virus: AAV2/9-hSyn-DIO-ChrimsonR-mCherry-WPRE-pA. Error bar, s.e.m.

(C) Significant spatial correlations of axon distribution of ChAT neurons expressed with optogenetic protein and fluorescence protein. Each dot represented an individual brain region in Figure A-B, scaled according to the voxel number of each region. C.C. Pearson's correlation coefficients.

(D) No significant difference of the spatial similarities between functional activation maps and anatomical projections with different virus expressed in ChAT-Cre mice. Each dot represented an individual EPI run. Error bar, ± s.t.d.

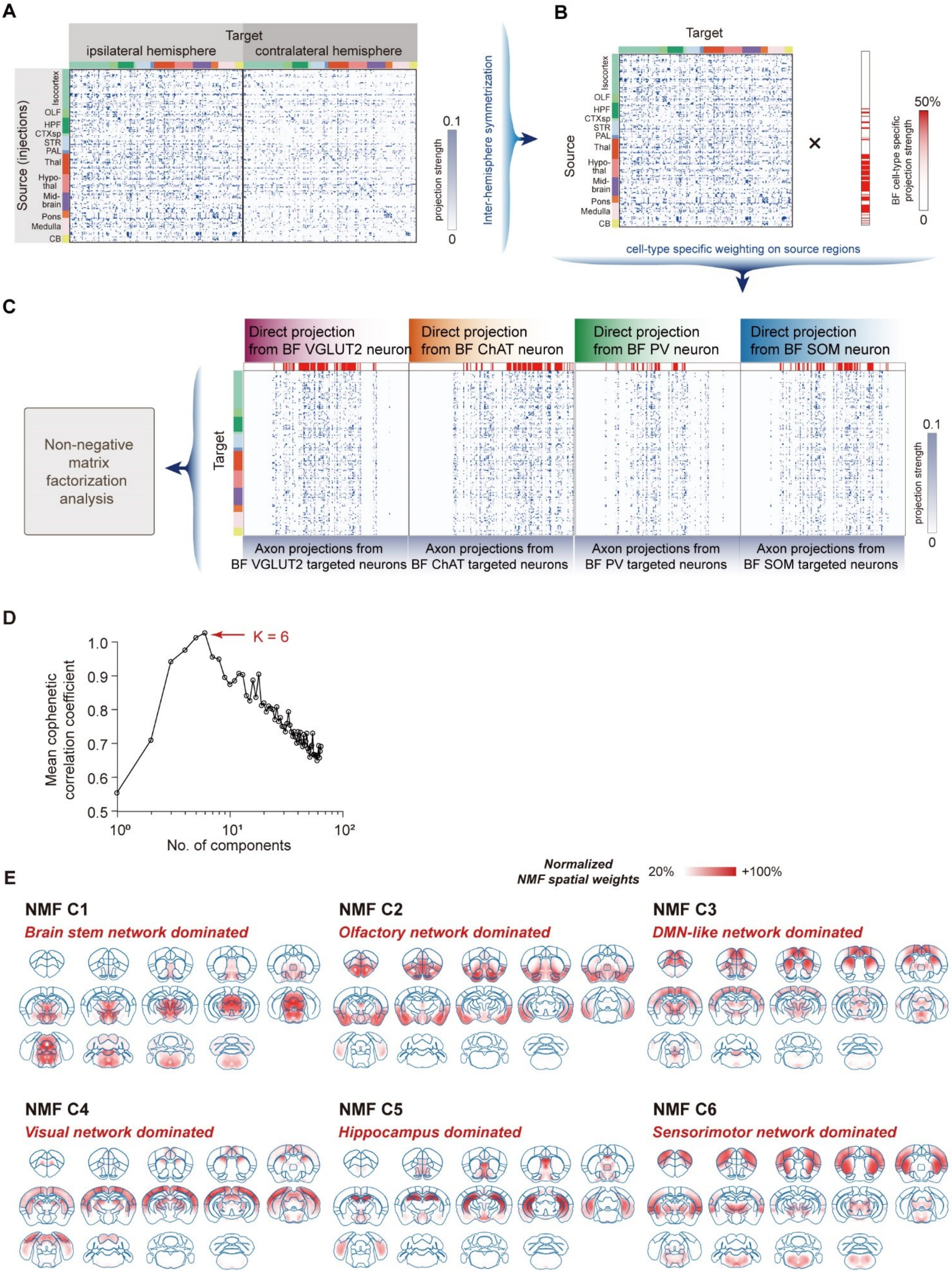


*(Legend on next page)*

**Supplementary Figure 10 | Related to Main Figure 4**

**Basal forebrain (BF) originated low dimensional structural networks.**

***(A-C) Computational pipeline of non-negative matrix factorization (NMF) on concatenated whole-brain mesoscopic secondary projections.***

(A) The whole-brain mesoscopic structural connectivity matrix was derived from the Voronoi resampled structural connectome in previous studies[3,4].

(B) With regard to the bilateral optogenetic stimulation of BF neurons, we computed the average connectivity strengths from both ipsilateral and contralateral hemispheres in order to construct a mesoscopic whole-brain structural connectivity matrix.

(C) To further investigate BF modulation of global functional networks, we utilized whole-brain mesoscopic structural connectivity weighted by each cell-type output from BF neurons. Subsequently, NMF was performed on the resulting connectivity matrix.

***(D-E) Top 6 non-negative factorization (NMF) components of the whole-brain mesoscopic connectome.***

(D) Mean cophenetic correlation coefficients were calculated for various numbers of NMF components, with the highest coefficient observed at k = 6, indicating that the top six NMF components represented the optimal subsets of the whole-brain mesoscopic connectome.

(E) Spatial weights of top 6 NMF components shown in the axial slice view. C[#], Component (e.g. C1, C2, C3).

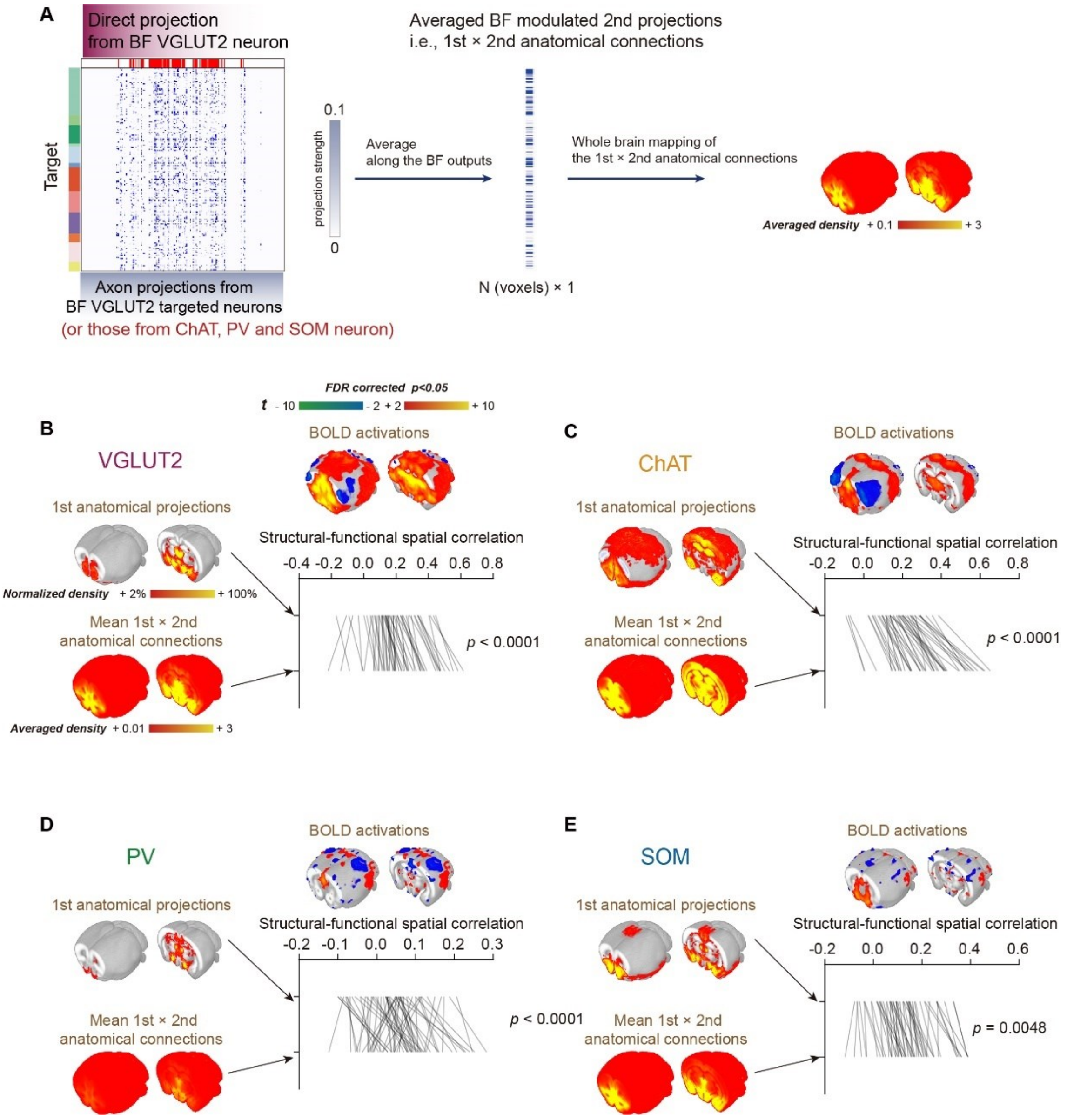


**Supplementary Figure 11 | Related to Main Figure 4**

**Two-synaptic connections showed higher spatial similarity of cell-type specific opto-fMRI activations.**

(A) Computational pipeline of the averaged 1st × 2nd anatomical connections. The weighted anatomical connection matrix (as in Figure S12C) was calculated through multiplying the Allen Mouse Brain mesoscopic connectivity matrix (i.e., 2nd connections) by each cell-type output from BF neurons (i.e., 1st projections) on the source direction. Then, the strength of 1st × 2nd anatomical connection matrix was averaged along the source direction, resulting the averaged 1st × 2nd anatomical connections.

(B) Higher structural-functional spatial correlation between BOLD activation map and averaged 1st × 2nd anatomical connections for VGLUT2 group. The spatial map of 1st anatomical projections was same as Main Figure 3B. Each line indicated an individual EPI run. Statistical significance was calculated using the two-tailed paired t-test.

(C-E) As in (B) but for BF ChAT, PV and SOM neurons.

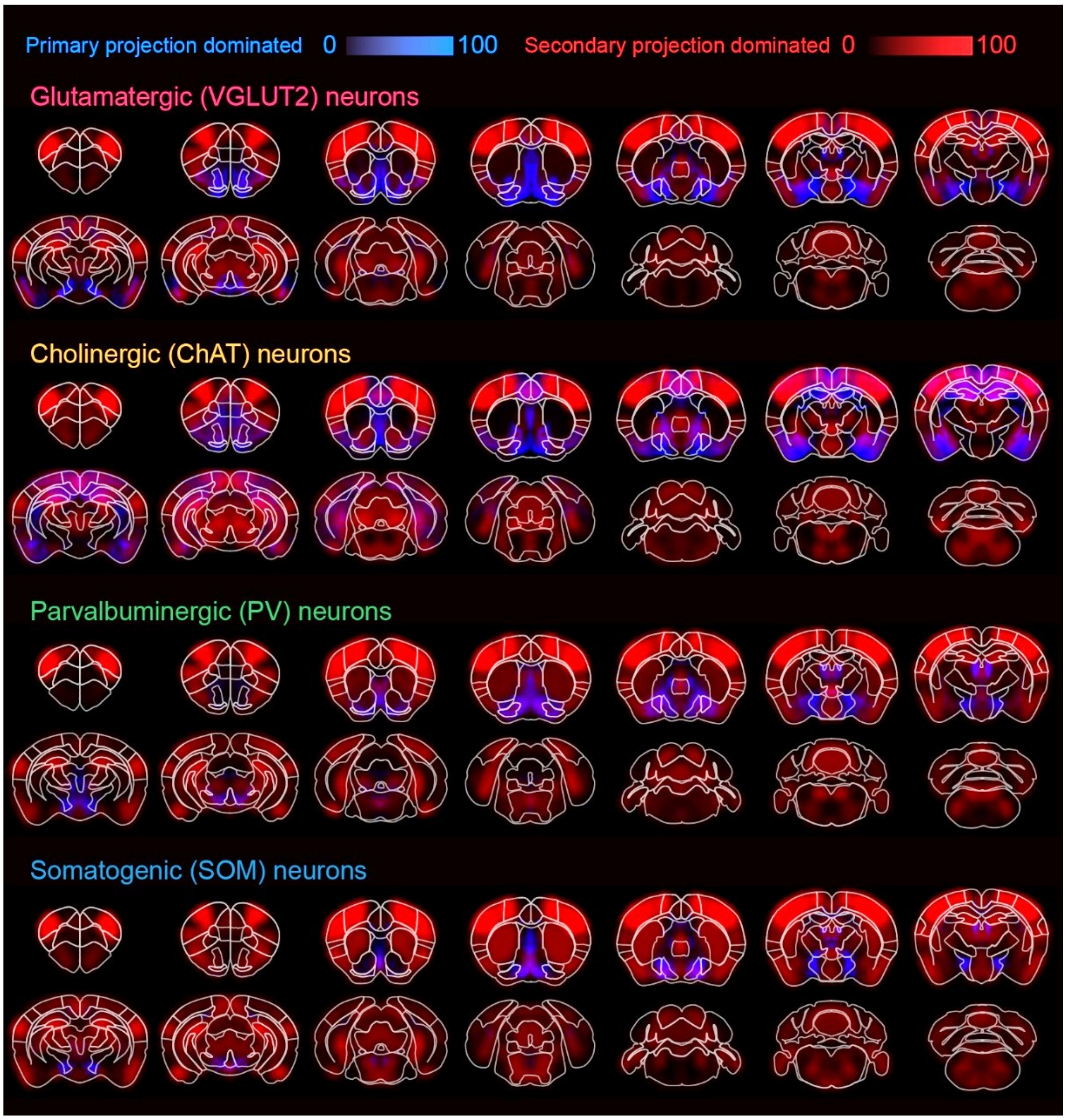


**Supplementary Figure 12 | Related to Main Figure 4**
**Spatial distributions of explained variance for opto-fMRI activations from the direct projections and low dimensional structural networks.**
Same as the Main Figure 4E, but shown in the axial slice view.

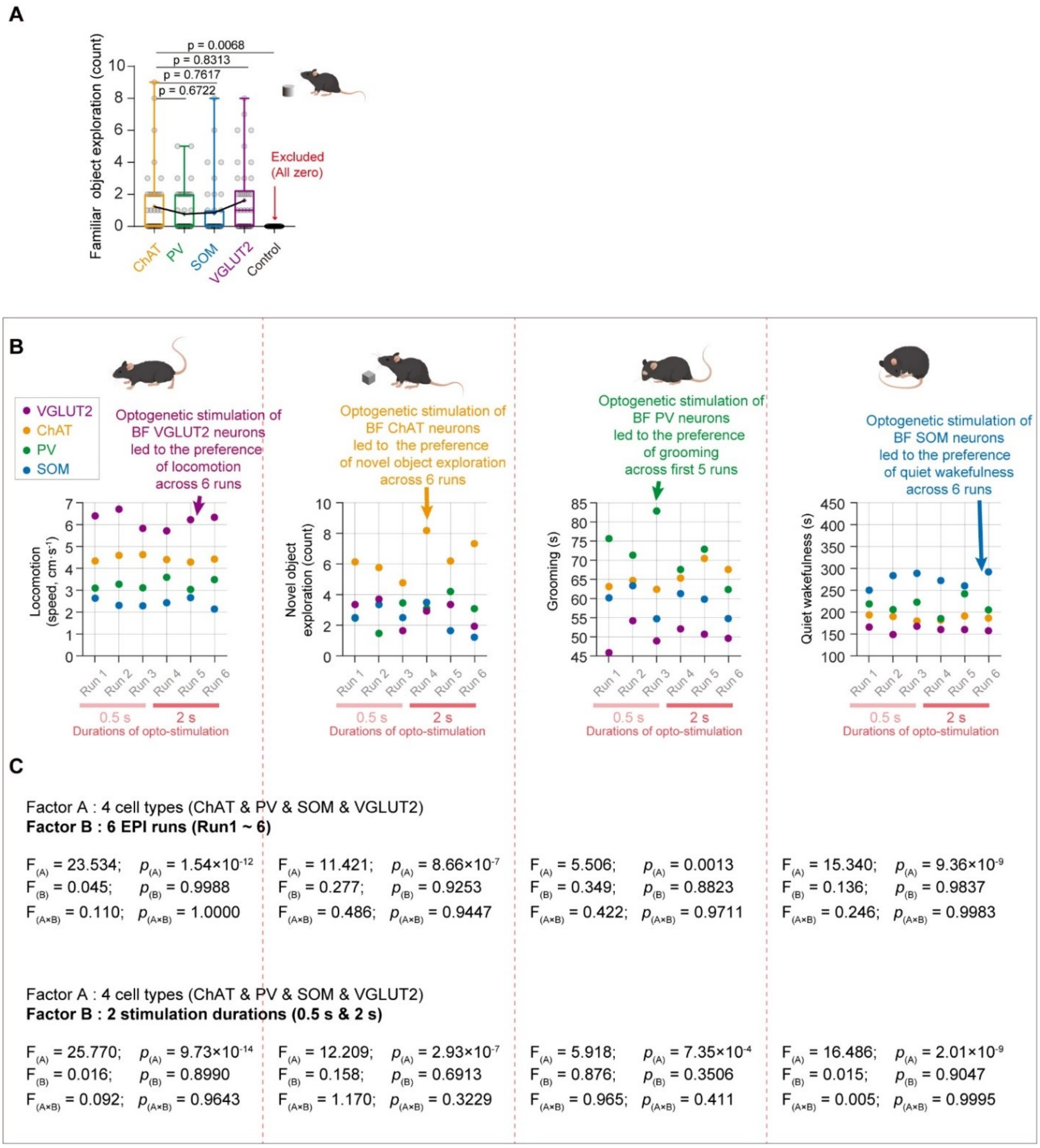


**Supplementary Figure 13 | Related to Main Figure 5**

**Significant contribution of BF neuron types to mouse behavioral preference across six runs.**

(A) Familiar object exploration was conducted under optogenetic stimulation of cell-type specific BF neurons, and no significant differences were observed among the groups. Each dot represented the averaged behavioral performance in each experimental session. Box, 25-75% range and median line; whisker, minimum to maximum range; plus mark, the mean.

(B) Behavioral preference modulated by BF neuron types remained consistent throughout all six runs. Behavioral performances under optogenetic activations of four BF neuron types, including locomotion, novel object exploration, grooming and quiet wakefulness. Colored dots represented averaged behavioral performance across mice.

(C) Two-way ANOVA analysis showed BF cell types dominated mouse behavioral preference and no significant effect on the mouse behaviors from experimental runs or durations.

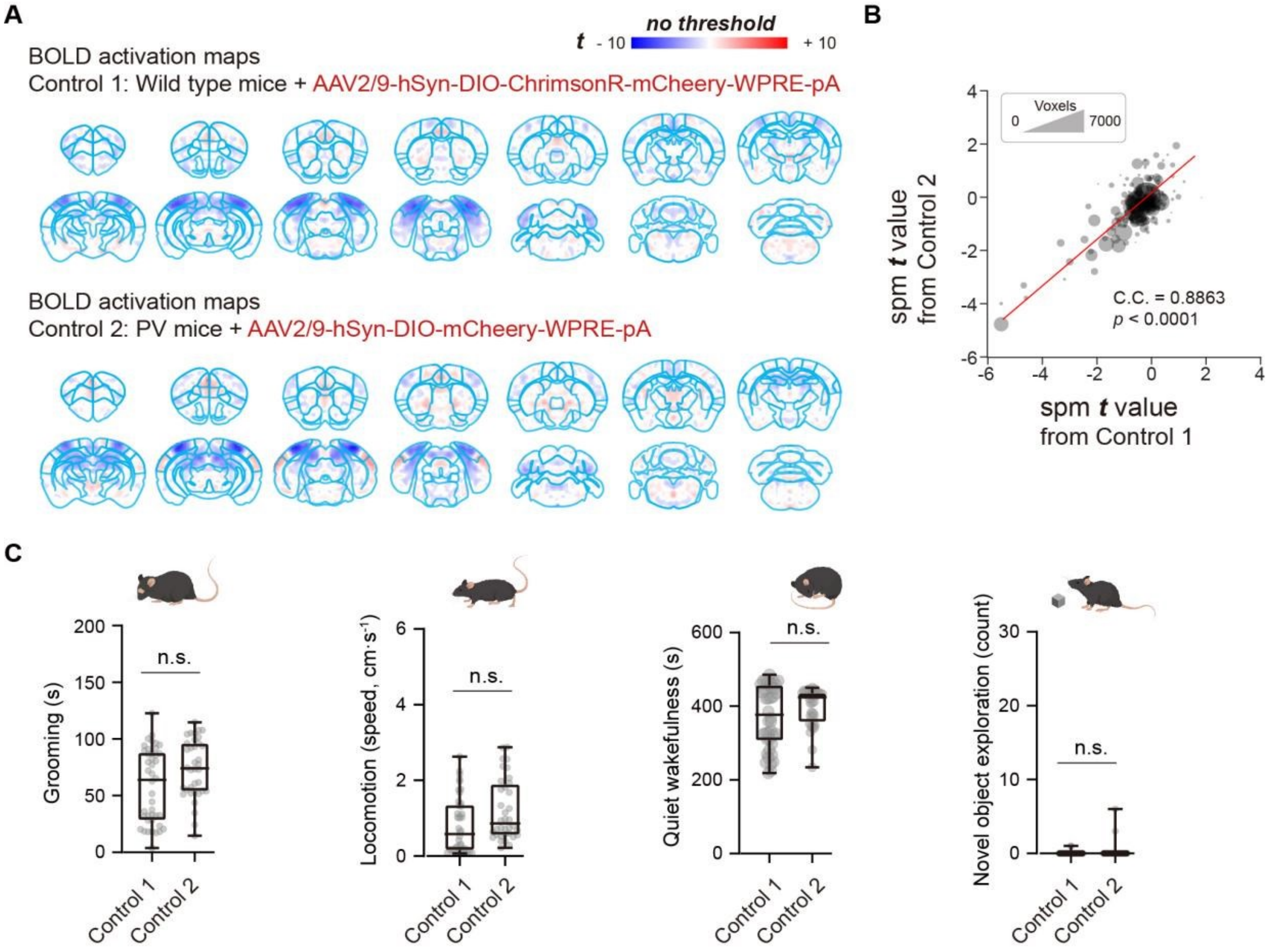


**Supplementary Figure 14 | Related to Main Figure 5**

**No significant difference of BOLD responses and mouse behaviors from two different control groups.**

(A) BOLD activation maps of control groups under optogenetic stimulations of BF neurons from two different control groups. BOLD activation maps were generated using the one-sample t-test.

(B) Significant spatial correlations between the BOLD activation maps from two control groups. Each dot represented an individual brain region, scaled according to the voxel number of each region. Red line represented the best linear fit. C.C. Pearson's correlation coefficients.

(C) No significant difference of mouse behaviors from two different animal cohorts. Statistical significance was determined using the two-sample t-test. Each dot represented an individual experimental session. Box, 25-75% range and median line; whisker, minimum to maximum range.

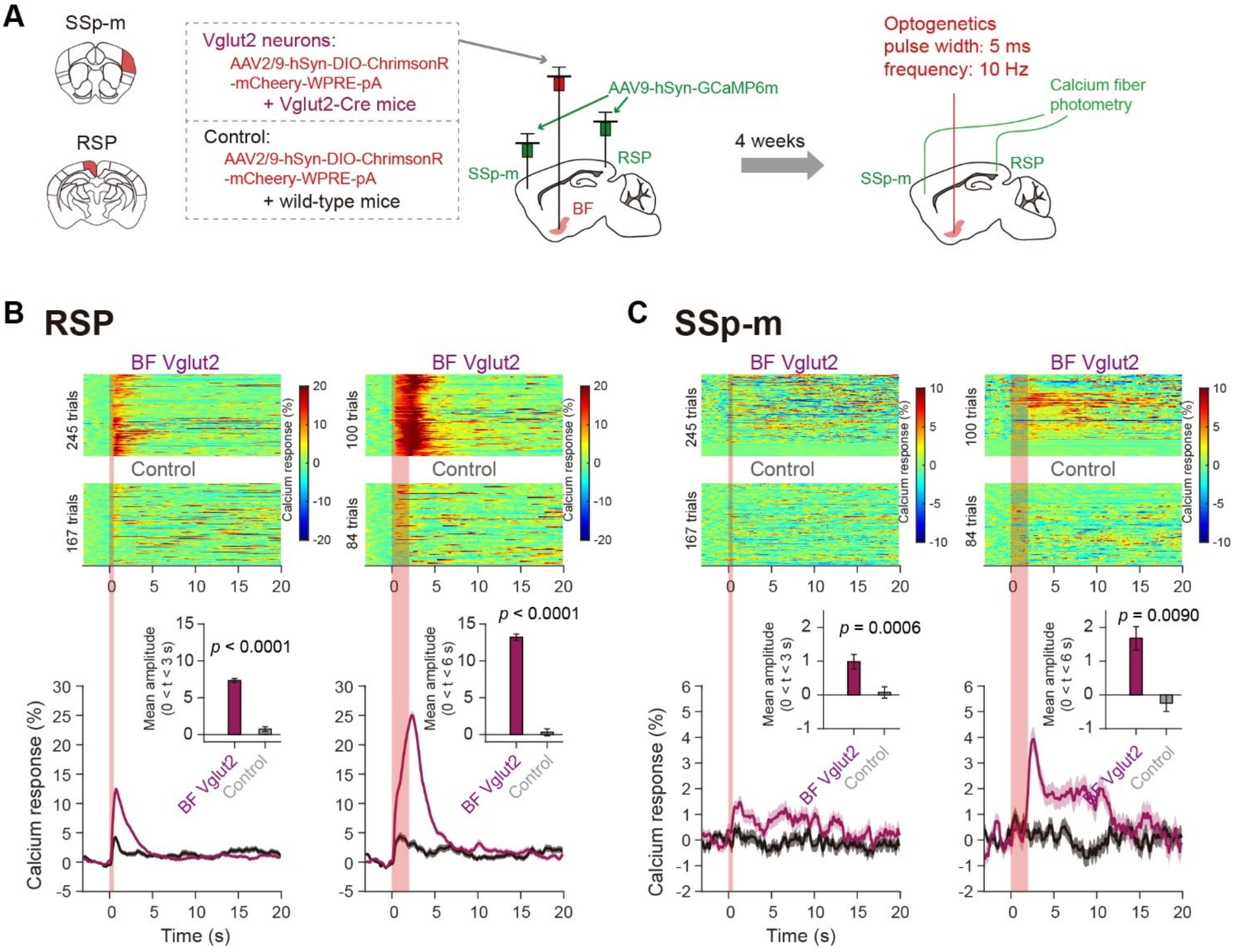


**Supplementary Figure 15 | Related to Main Figure 5**

**The initial rise of BOLD signals correlated to the increased calcium activities evoked by optogenetics in RSP and SSp-m.**

(A) Schematic diagram depicting fiber photometry recording of neuronal calcium signals in RSP and SSp-m under the optogenetic stimulation on BF neurons for VGLUT2 or Control group. AAVs expressing DIO-ChrimsonR-mCheery or DIO-mCheery were injected into the bilateral BF, and AAVs expressing GCaMP6m were injected into the unilateral SSp-m and RSP. SSp-m, primary somatosensory area (mouth); RSP, retrosplenial area.

(B) Optogenetic activations of BF VGLUT2 neurons evoked significant elevations of neuronal activities in RSP. Upper, optogenetic stimulation evoked calcium signal changes in RSP across different trials ($N_{VGLUT2}$ = 5 mice and $N_{control}$ = 4 mice). Lower, as in (Upper) but for the averaged calcium responses. Error bars represented the standard error of the mean (SEM). Lower insert, quantitative evaluation of calcium signal amplitude in RSP between VGLUT2 and Control groups. Statistical significance was determined using the two-tailed two sample t-test. Error bar, standard error of the mean (SEM).

(C) As in (B) but for the calcium signals in SSp-m. Notably, neuronal activities in SSp-m evoked by BF VGLUT2 neurons were significantly higher than those in Control group as well.

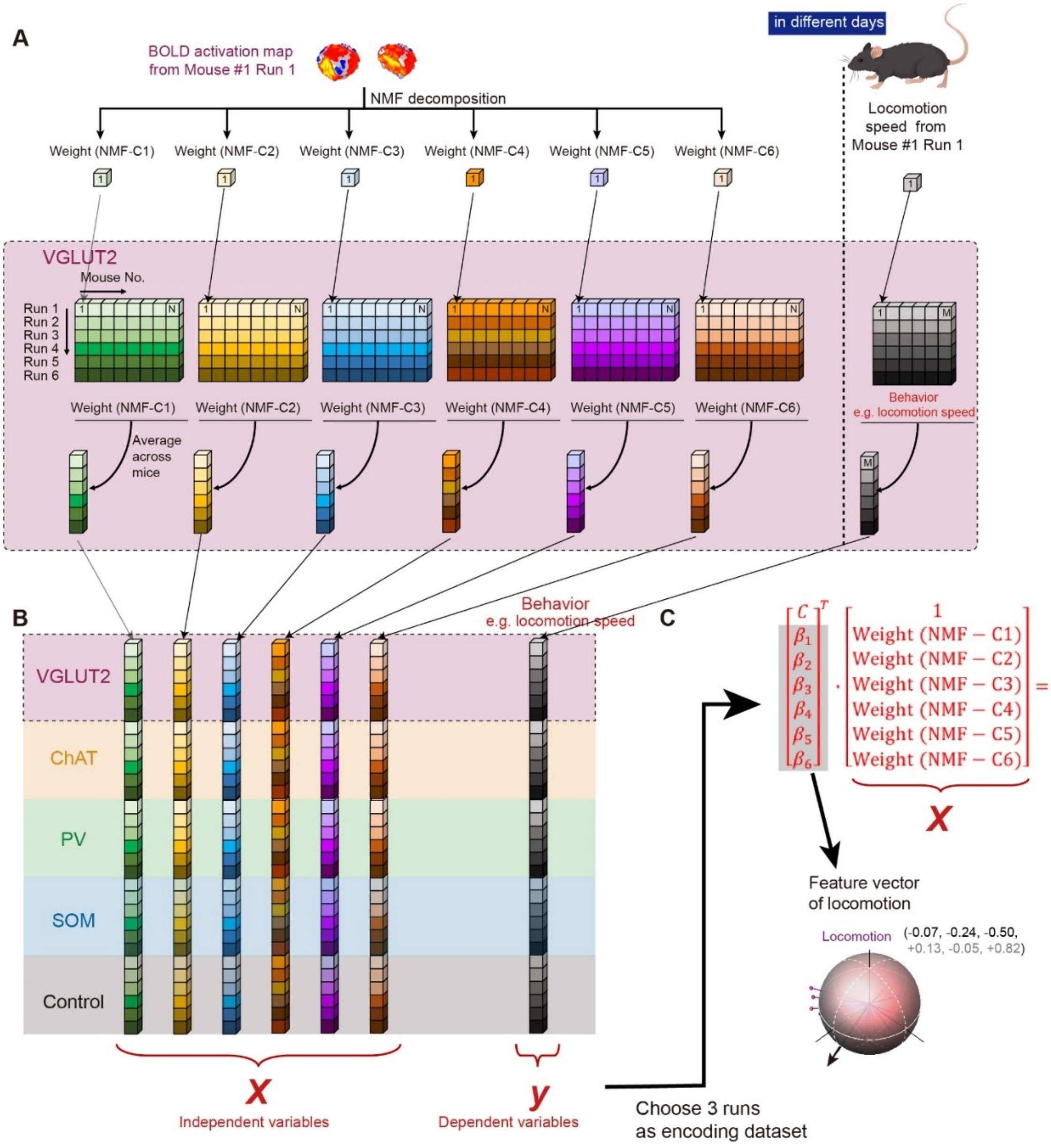


**Supplementary Figure 16 | Related to Main Figure 6**

**Computational pipeline for the decoding model to link behaviors and BF modulated BOLD activations (<u>took the locomotion behavior as the example</u>).**

(A) As the optogenetic fMRI and free-moving behavioral tests were conducted in different days, we divided the total behavior length into 6 runs for each mouse, keeping temporal alignment with the order of fMRI scanning. And also, the BOLD activation maps were decomposed by the NMF strategy. Then, we averaged both the weights of NMF components and behavioral performances across mice, and only considered the group-wise behaviors under optogenetic stimulations on cell-type specific neurons.

(B) The resulting cell-type specific BOLD weights of NMF components ($W_{5\times6\times6}$) were set as the model predictors (5 groups, 6 runs and 6 NMF components), and the behavioral performances ($Y_{5\times6}$) were termed as the model output.

(C) Regression coefficients ($\beta_0$, $\beta_1$, $\beta_2$, $\beta_3$, $\beta_4$, $\beta_5$ and $\beta_6$) were termed as the representative vector $[\beta_1, \beta_2, \beta_3, \beta_4, \beta_5, \beta_6]$ for each mouse behavior in NMF space, and further used to construct corresponding spatial patterns by multiplying the spatial NMF components.

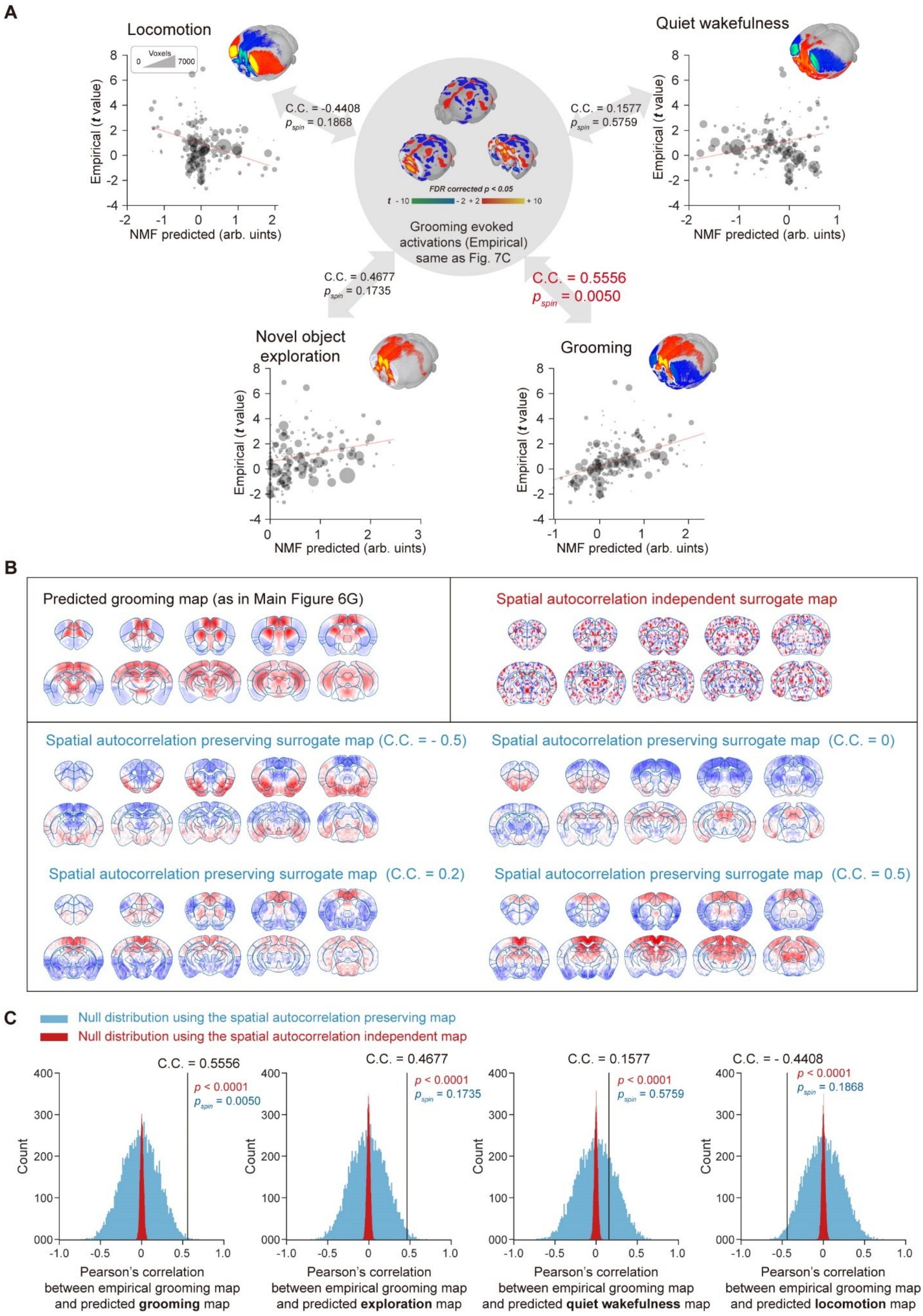


*(Legend on next page)*

**Supplementary Figure 17 | Related to Main Figure 7**

**Spatial autocorrelation preserving surrogate maps provided a more conservative and meaningful measure of statistical significance for the similarity of spatial maps.**

(A) Significant spatial correlation between empirical and predicted grooming maps. Null distribution was derived from the Pearson's correlation coefficients (C.C.) between empirical grooming map and spatial autocorrelation preserving surrogate maps of each predicted behavioral spatial map. Each dot represented an individual brain region, scaled according to the voxel number of each region. C.C. Pearson's correlation coefficients.

(B) The predicted spatial pattern of mouse grooming and representative surrogate maps with matched independent and preserving spatial autocorrelation.

(C) Null distribution of Pearson's correlations between empirical grooming map and predicted maps of mouse behaviors (sky blue, 10000 shuffled spatial autocorrelations preserving; red, 10000 shuffled spatial autocorrelation-independent). Black lines represented the correlations between empirical grooming map and predicted maps of mouse behaviors.

**Supplementary Table 1 | Related to Main Figure 1**
**Abbreviations of mouse brain regions.**

| Abbreviation | Full Name | Major Regions |
|---|---|---|
| BF | Basal forebrain | Pallidum |
| Mot | Motor area | Isocortex |
| SS | Somatosensory area | Isocortex |
| AUD | Auditory area | Isocortex |
| VIS | Visual area | Isocortex |
| ORB | Orbital area | Isocortex |
| AI | Agranular insular area | Isocortex |
| ACA | Anterior cingulate area | Isocortex |
| RSP | Retrosplenial area | Isocortex |
| HPF | Hippocampal Formation | Hippocampal Formation |
| CA1 | Field CA1 | Hippocampal Formation |
| CA2 | Field CA2 | Hippocampal Formation |
| CA3 | Field CA3 | Hippocampal Formation |
| CP | Caudoputamen | Striatum |
| ACB | Nucleus accumbens | Striatum |
| DORsm | Thalamus, sensory-motor cortex related | Thalamus |
| DORpm | Thalamus, polymodal association cortex related | Thalamus |
| LZ | Hypothalamic lateral zone | Hypothalamus |
| MBsen | Midbrain, sensory related | Midbrain |
| MBmot | Midbrain, motor related | Midbrain |
| MBsta | Midbrain, behavioral state related | Midbrain |
| Pons | / | Pons |
| Medulla | / | Medulla |
| CBX | Cerebellar cortex | Cerebellum |

**Supplementary Table 2 | Multiple Comparisons of mouse behavioral performance under optogenetic stimulations of four BF cell types.**

(a) locomotion

| ANOVA table | SS (Type III) | DF | MS | F (DFn, DFd) | P value |
|---|---|---|---|---|---|
| Interaction | 9.366 | 20 | 0.4683 | F (20, 186) = 0.1149 | P>0.9999 |
| Sessions | 1.661 | 5 | 0.3322 | F (5, 186) = 0.08154 | P=0.9950 |
| BF cell types | 692.6 | 4 | 173.1 | F (4, 186) = 42.50 | P<0.0001 |
| Residual | 757.8 | 186 | 4.074 | | |

| Tukey's multiple comparisons test | mean diff. | 95.00% CI of diff. | Significant? | Summary | Adjusted P Value |
|---|---|---|---|---|---|
| VGLUT2 vs. ChAT | 1.765 | 0.5512 to 2.978 | Yes | *** | 0.0008 |
| VGLUT2 vs. PV | 2.950 | 1.775 to 4.125 | Yes | **** | <0.0001 |
| VGLUT2 vs. SOM | 3.810 | 2.597 to 5.024 | Yes | **** | <0.0001 |
| VGLUT2 vs. Ctrl | 5.356 | 4.143 to 6.569 | Yes | **** | <0.0001 |
| ChAT vs. PV | 1.185 | 0.01041 to 2.360 | Yes | * | 0.0468 |
| ChAT vs. SOM | 2.046 | 0.8325 to 3.259 | Yes | **** | <0.0001 |
| ChAT vs. Ctrl | 3.591 | 2.378 to 4.805 | Yes | **** | <0.0001 |
| PV vs. SOM | 0.8606 | -0.3143 to 2.035 | No | ns | 0.2615 |
| PV vs. Ctrl | 2.406 | 1.231 to 3.581 | Yes | **** | <0.0001 |
| SOM vs. Ctrl | 1.546 | 0.3322 to 2.759 | Yes | ** | 0.0051 |

(b) novel object exploration

| ANOVA table | SS (Type III) | DF | MS | F (DFn, DFd) | P value |
|---|---|---|---|---|---|
| Interaction | 144.5 | 20 | 7.225 | F (20, 186) = 0.4150 | P=0.9879 |
| Sessions | 28.87 | 5 | 5.774 | F (5, 186) = 0.3317 | P=0.8933 |
| BF cell types | 723.6 | 4 | 180.9 | F (4, 186) = 10.39 | P<0.0001 |
| Residual | 3238 | 186 | 17.41 | | |

| Tukey's multiple comparisons test | mean diff. | 95.00% CI of diff. | Significant? | Summary | Adjusted P Value |
|---|---|---|---|---|---|
| VGLUT2 vs. ChAT | -3.250 | -5.758 to -0.7420 | Yes | ** | 0.0041 |
| VGLUT2 vs. PV | -0.1429 | -2.571 to 2.286 | No | ns | 0.9998 |
| VGLUT2 vs. SOM | 0.3690 | -2.139 to 2.877 | No | ns | 0.9943 |
| VGLUT2 vs. Ctrl | 2.560 | 0.05149 to 5.068 | Yes | * | 0.0429 |
| ChAT vs. PV | 3.107 | 0.6787 to 5.536 | Yes | ** | 0.0048 |
| ChAT vs. SOM | 3.619 | 1.111 to 6.127 | Yes | *** | 0.0009 |
| ChAT vs. Ctrl | 5.810 | 3.301 to 8.318 | Yes | **** | <0.0001 |
| PV vs. SOM | 0.5119 | -1.916 to 2.940 | No | ns | 0.9778 |
| PV vs. Ctrl | 2.702 | 0.2740 to 5.131 | Yes | * | 0.0208 |
| SOM vs. Ctrl | 2.190 | -0.3176 to 4.699 | No | ns | 0.1183 |

### (c) quiet wakefulness

| ANOVA table | SS (Type III) | DF | MS | F (DFn, DFd) | P value |
|---|---|---|---|---|---|
| Interaction | 39754 | 20 | 1988 | F (20, 186) = 0.2900 | P=0.9989 |
| Sessions | 5769 | 5 | 1154 | F (5, 186) = 0.1683 | P=0.9740 |
| BF cell types | 1214973 | 4 | 303743 | F (4, 186) = 44.32 | P<0.0001 |
| Residual | 1274802 | 186 | 6854 | | |

| Tukey's multiple comparisons test | mean diff. | 95.00% CI of diff. | Significant ? | Summary | Adjusted P Value |
|---|---|---|---|---|---|
| VGLUT2 vs. ChAT | -27.26 | -77.02 to 22.51 | No | ns | 0.5581 |
| VGLUT2 vs. PV | -54.34 | -102.5 to -6.154 | Yes | * | 0.0184 |
| VGLUT2 vs. SOM | -115.1 | -164.8 to -65.29 | Yes | **** | <0.0001 |
| VGLUT2 vs. Ctrl | -213.3 | -263.1 to -163.5 | Yes | **** | <0.0001 |
| ChAT vs. PV | -27.08 | -75.27 to 21.10 | No | ns | 0.5326 |
| ChAT vs. SOM | -87.80 | -137.6 to -38.03 | Yes | **** | <0.0001 |
| ChAT vs. Ctrl | -186.1 | -235.8 to -136.3 | Yes | **** | <0.0001 |
| PV vs. SOM | -60.72 | -108.9 to -12.53 | Yes | ** | 0.0058 |
| PV vs. Ctrl | -159.0 | -207.2 to -110.8 | Yes | **** | <0.0001 |
| SOM vs. Ctrl | -98.26 | -148.0 to -48.49 | Yes | **** | <0.0001 |

### (d) grooming

| ANOVA table | SS (Type III) | DF | MS | F (DFn, DFd) | P value |
|---|---|---|---|---|---|
| Interaction | 3035 | 20 | 151.7 | F (20, 186) = 0.2985 | P=0.9987 |
| Sessions | 655.2 | 5 | 131.0 | F (5, 186) = 0.2578 | P=0.9355 |
| BF cell types | 12287 | 4 | 3072 | F (4, 186) = 6.043 | P=0.0001 |
| Residual | 94544 | 186 | 508.3 | | |

| Tukey's multiple comparisons test | mean diff. | 95.00% CI of diff. | Significant ? | Summary | Adjusted P Value |
|---|---|---|---|---|---|
| VGLUT2 vs. ChAT | -15.47 | -29.02 to -1.915 | Yes | * | 0.0164 |
| VGLUT2 vs. PV | -22.00 | -35.12 to -8.880 | Yes | **** | <0.0001 |
| VGLUT2 vs. SOM | -8.841 | -22.39 to 4.712 | No | ns | 0.3784 |
| VGLUT2 vs. Ctrl | -8.044 | -21.60 to 5.509 | No | ns | 0.4771 |
| ChAT vs. PV | -6.534 | -19.66 to 6.588 | No | ns | 0.6465 |
| ChAT vs. SOM | 6.627 | -6.926 to 20.18 | No | ns | 0.6622 |
| ChAT vs. Ctrl | 7.425 | -6.128 to 20.98 | No | ns | 0.5579 |
| PV vs. SOM | 13.16 | 0.03877 to 26.28 | Yes | * | 0.0489 |
| PV vs. Ctrl | 13.96 | 0.8364 to 27.08 | Yes | * | 0.0309 |
| SOM vs. Ctrl | 0.7976 | -12.76 to 14.35 | No | ns | 0.9998 |
